# Genome-wide strategies identify molecular niches regulated by connective tissue-associated transcription factors

**DOI:** 10.1101/165837

**Authors:** Mickael Orgeur, Marvin Martens, Georgeta Leonte, Sonya Nassari, Marie-Ange Bonnin, Stefan T. Börno, Bernd Timmermann, Jochen Hecht, Delphine Duprez, Sigmar Stricker

## Abstract

**Background:** Connective tissues support, connect and separate tissues and organs, playing crucial roles in development, homeostasis and fibrosis. Cell specification and differentiation is triggered by the activity of specific transcription factors. While key transcription factors have been identified for differentiation processes of most tissues, connective tissue differentiation remains largely unstudied.

**Results:** To gain insight into the regulatory cascades involved in connective tissue differentiation, we selected five zinc finger transcription factors - OSR1, OSR2, EGR1, KLF2 and KLF4 - based on their expression patterns and/or known involvement in the differentiation of mesenchymal cells into connective tissue subtypes. We combined RNA-seq with ChIP-seq profiling in chick limb cells following overexpression of individual transcription factors. We identified a set of common genes regulated by all five transcription factors, which constitutes a connective tissue core expression set. This common core was enriched in genes associated with axon guidance and myofibroblast signature. In addition, each of the transcription factors regulated a different set of extracellular matrix components and signalling molecules, which define local molecular niches important for connective tissue development and function.

**Conclusions:** The established regulatory network identifies common and distinct molecular signatures downstream of five connective tissue-associated transcription factors and provides insight into the signalling pathways governing limb connective tissue differentiation. It also suggests a concept whereby local molecular niches can be created via the expression of specific transcription factors impinging on the specification of microenvironments.

## Background

All cells in a multicellular eukaryotic organism share the same genetic information. However, the choice of specification, differentiation and function taken by a progenitor cell encompasses dramatic transcriptional and finally phenotypic changes that differ for each cell/tissue type. Lineage-specific genetic programs consist in a fine-tuning between the repression and the expression of a given set of genes in response to extrinsic and intrinsic signals at a specific location and/or at a precise time [1]. Progenitor cells are specified and induced to differentiate along a certain lineage upon activation of lineage-specific key transcription factors (TFs) that drive specific transcriptional programs [2]. As a consequence, the differentiating progenitor cells express lineage-specific genes that reinforce lineage commitment, as well as providing unique characteristics to the specific cell type and the tissue it gives rise to.

Connective tissue (CT) is one of the main components of the body supporting tissues and organs. The term of CT gathers together an ensemble of tissues such as specialized CT (cartilage and bone), soft CT (adipose tissue and vasculature) and dense CT. Dense CT can be divided into regular CT (tendon and ligament) and irregular CT (loose CT surrounding or within organs such as muscle CT) [3]. Regular and irregular CTs are part of the musculoskeletal system and are associated with cartilage/bone and skeletal muscle. Dysregulation of CT homeostasis leads to fibrosis, which is observed during pathological tissue repair or healing processes and in cancer [4]. Although fibrosis is a common research subject, normal CT formation during development remains to date poorly investigated.

The appendage of vertebrate embryos is an excellent model system for analysing tissue differentiation and cellular interactions during development. In limbs, cells forming the skeleton, as well as regular and irregular CTs, are derived from the lateral plate mesoderm, while myogenic cells originate from the somites [5,6]. Classical embryological experiments have shown that limb patterning is dependent on lateral plate-derived CTs that provide instructive cues to guarantee correct muscle, nerve and vessel formation [7–10]. This indicates that CT cells are key players creating local microenvironments that contain permissive and/or instructive cues for organ patterning. While master TFs governing cell-type specific gene expression programs have been identified for cartilage (SOX5, SOX6, SOX9), bone (OSX, RUNX2) and muscle (MYF5, MYF6, MYOD) development [11,12], knowledge is sparse for dense CT. CT is mostly identified by gene/protein expression associated with CT function. Irregular CT is associated with type-III and -VI collagens, while regular CT (tendon/ligament) is characterized by the expression of structural and functional components such as type-I and -XII collagens and Tenomodulin (TNMD) [7,13]. Few TFs specific for CT lineages have been identified. Scleraxis (SCX) is to date the unique marker for tendon progenitors, however it is not necessary for development of most tendons [14,15]. Early growth response 1 (EGR1) is also not required for mouse tendon development but is involved in type-I collagen production in chick and mouse developing tendons [16]. Moreover, EGR1 forced expression is sufficient to induce the expression of tendon-associated genes in murine mesenchymal stem cells [17]. Limb irregular CT associated with muscle is marked by the expression of TCF4 (TCF7l2), TBX4 or TBX5, but they have no obvious role in CT differentiation [18,19]. In contrast, Odd-skipped related 1 and 2 (OSR1 and OSR2) are expressed and involved in loose irregular CT differentiation during chick and mouse limb development [20,21,Vallecillo Garcia et al. in revision].

Here, we analysed the molecular mechanisms underlying CT differentiation and function during chick limb development, in order to provide a framework for future analyses of CT development and CT-muscle interconnectivity. To this end, we selected five zinc finger TFs involved or presumably involved in CT differentiation. OSR1, OSR2 and EGR1 were chosen based on their demonstrated contribution in irregular and regular CT development, respectively. KLF2 and KLF4 (Krüppel-like factor 2 and 4) were chosen based on their expression patterns in CT associated with tendons, although their role in limb development is presently not elucidated. We combined whole-transcriptome sequencing (RNA-seq) and chromatin immunoprecipitation followed by massively parallel DNA sequencing (ChIP-seq) to identify the gene regulatory programs downstream of each TF. This allowed us to design a novel, unique and unexplored global regulatory network underlying CT differentiation and to identify common and specific molecular niches that are shaped during this process.

## Results

### Limb expression patterns of CT-associated TFs

We investigated gene expression patterns of the five selected CT-associated TFs in chick limbs during development. Consistent with previous observations [20,21], *OSR1* and *OSR2* were expressed in dorsal and ventral limb regions of E4.5 chick embryos (Fig. 1a, b). *OSR1* and *OSR2* expression domains overlapped with those of *SCX* and *MYOD*, which labelled tendon and myogenic cells, respectively (Fig. 1c, d). In contrast to both *OSR* transcripts, *EGR1*, *KLF2* and *KLF4* were not detected in E4.5 limb buds. At later stages of limb development, when the initial pattern of the musculoskeletal system is set, both *OSR1* and *OSR2* were not expressed in *SCX*-positive tendons, but rather expressed in muscle CT, interstitial to muscle fibres, while *OSR1* was also detected surrounding individual muscles (Fig. 1e-i). *EGR1* was expressed in tendons, close to muscle attachments (Fig 1j, k), as previously described [16], while *KLF2* and *KLF4* transcripts delineated *SCX*-positive tendons of the knee of E9.5 chick embryos (Fig. 1l-q). In summary, *OSR1* and *OSR2* label irregular CT, whereas *EGR1*, *KLF2* and *KLF4* are expressed in different regions of regular CT in developing chick limbs.

**Fig. 1.**
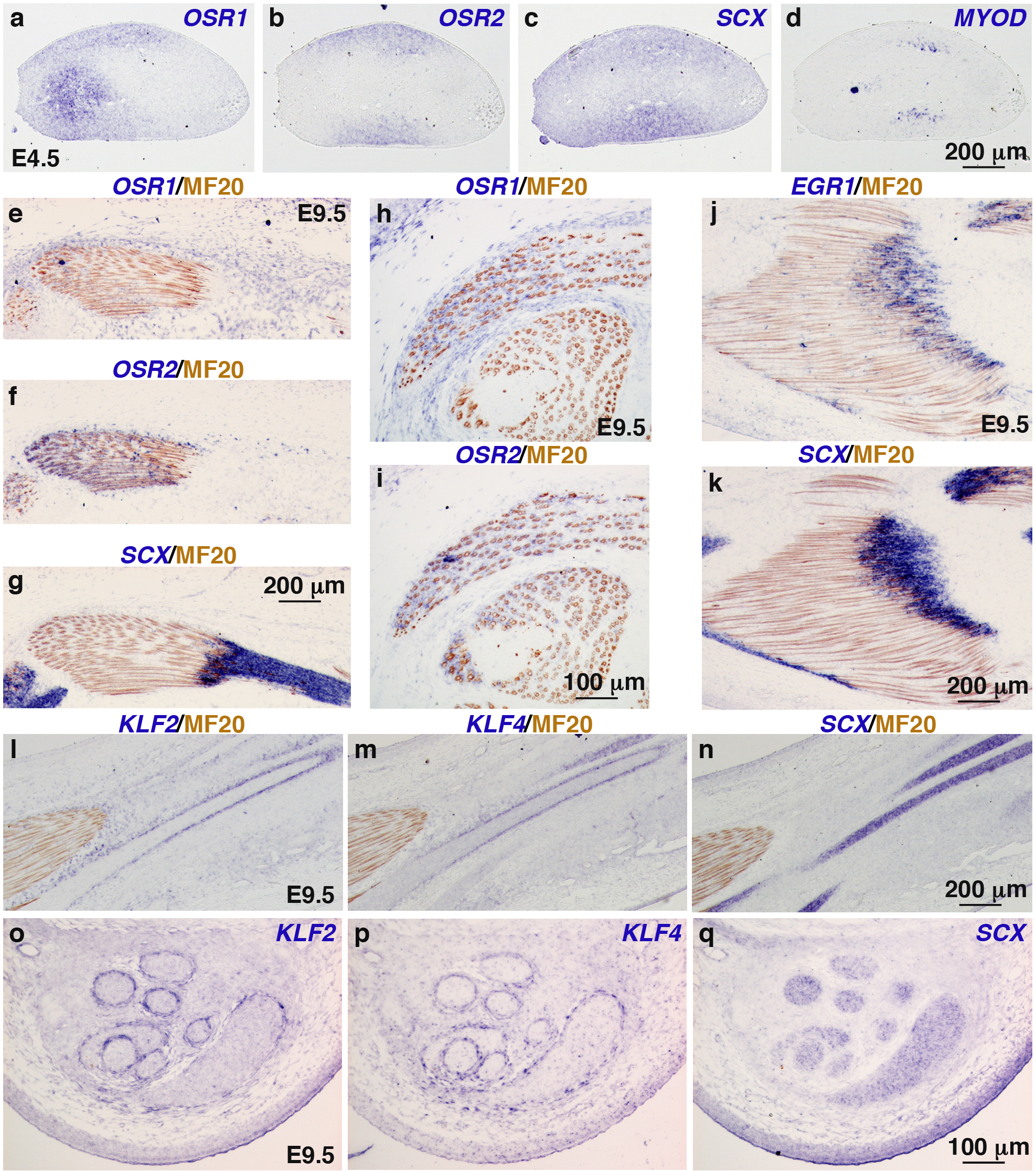
Endogenous expression of CT-associated TFs in hindlimbs of chick embryos. (**a**-**d**) In situ hybridization to hindlimbs of E4.5 chick embryos. Adjacent and transverse limb sections were hybridized with *OSR1* (**a**), *OSR2* (**b**), *SCX* (**c**) and *MYOD* (**d**) probes (blue). (**e**-**q**) In situ hybridization to hindlimbs of E9.5 chick embryos followed by immunohistochemistry with the MF20 antibody (brown), which recognizes skeletal muscle myosins. (**e**-**g**) Adjacent and longitudinal limb sections were hybridized with *OSR1* (**e**), *OSR2* (**f**) and *SCX* (**g**) probes (blue). (**h**, **i**) Adjacent and transverse limb sections were hybridized with *OSR1* (**h**) and *OSR2* (**i**) probes (blue). (**j**, **k**) Adjacent and longitudinal limb sections were hybridized with *EGR1* (**j**) and *SCX* (**k**) probes (blue). (**l**-**n**) Adjacent and longitudinal limb sections were hybridized with *KLF2* (**l**), *KLF4* (**m**) and *SCX* (**n**) probes (blue). (**o**-**q**) Adjacent and transverse limb sections were hybridized with *KLF2* (**o**), *KLF4* (**p**) and *SCX* (**q**) probes (blue).

### The selected TFs influence differentiation of limb mesenchymal cells

To analyse the functionality of the five TFs towards CT differentiation, we chose the chick micromass (chMM) explant model (Fig. 2a). In this three-dimensional culture model, primary limb bud cells behave close to the in vivo situation and differentiate into the mesenchymal lineages observed in native limb buds [22]. We tested the ability of the five TFs to shift cell differentiation via their retroviral overexpression in chMM cultures. While the overall morphology of the chMM cultures remained unchanged across all conditions, cartilage differentiation was affected upon TF overexpression (Fig. 2b, c). In agreement with previous observations [21], OSR1 and OSR2 overexpression reduced chondrogenic matrix production by 58% and 67%, respectively, compared to control cultures (Fig. 2c, d). Similarly, KLF2 and KLF4 overexpression induced a reduction of cartilage nodule formation, but to a lower extent compared to OSR1 and OSR2 (Fig. 2c, d). EGR1 was the only factor that increased chondrogenic matrix production within the chMM cultures (Fig. 2c, d). Quantitative RT-PCR analysis of transcript levels of cartilage-associated genes, *SOX9* and *COL2A1*, confirmed the inhibitory effect of OSR1, OSR2, KLF2 and KLF4 overexpression, as well as the positive effect of EGR1 overexpression on cartilage differentiation (Fig. 2e-g). EGR1 overexpression increased the expression of the tendon differentiation marker, *TNMD*, while not affecting that of the irregular CT markers *COL3A1* and *COL6A1* (Fig. 2e). Overexpression of KLF2, but not that of KLF4, increased the expression levels of the tendon markers *SCX* and *TNMD* (Fig. 2f). Both KLF factors also increased *COL6A1* expression (Fig. 2f). OSR2 overexpression increased the expression of the CT markers *COL3A1* and *COL6A1*, while OSR1 overexpression only affected *COL3A1* expression (Fig. 2g). In summary, the TFs had different outcome on CT differentiation. OSR1 and OSR2 drove undifferentiated limb mesenchymal cells towards irregular CT differentiation at the expense of cartilage differentiation. EGR1 induced tendon and cartilage marker expression, while not affecting irregular CT marker expression. KLF2, but not KLF4, promoted the expression of tendon markers, while both KLF factors increased *COL6A1* expression and decreased *COL2A1* expression. In conclusion, all five TFs proved functional in this model towards an effect on CT cell differentiation thus validating them for further analysis.

**Fig. 2.**
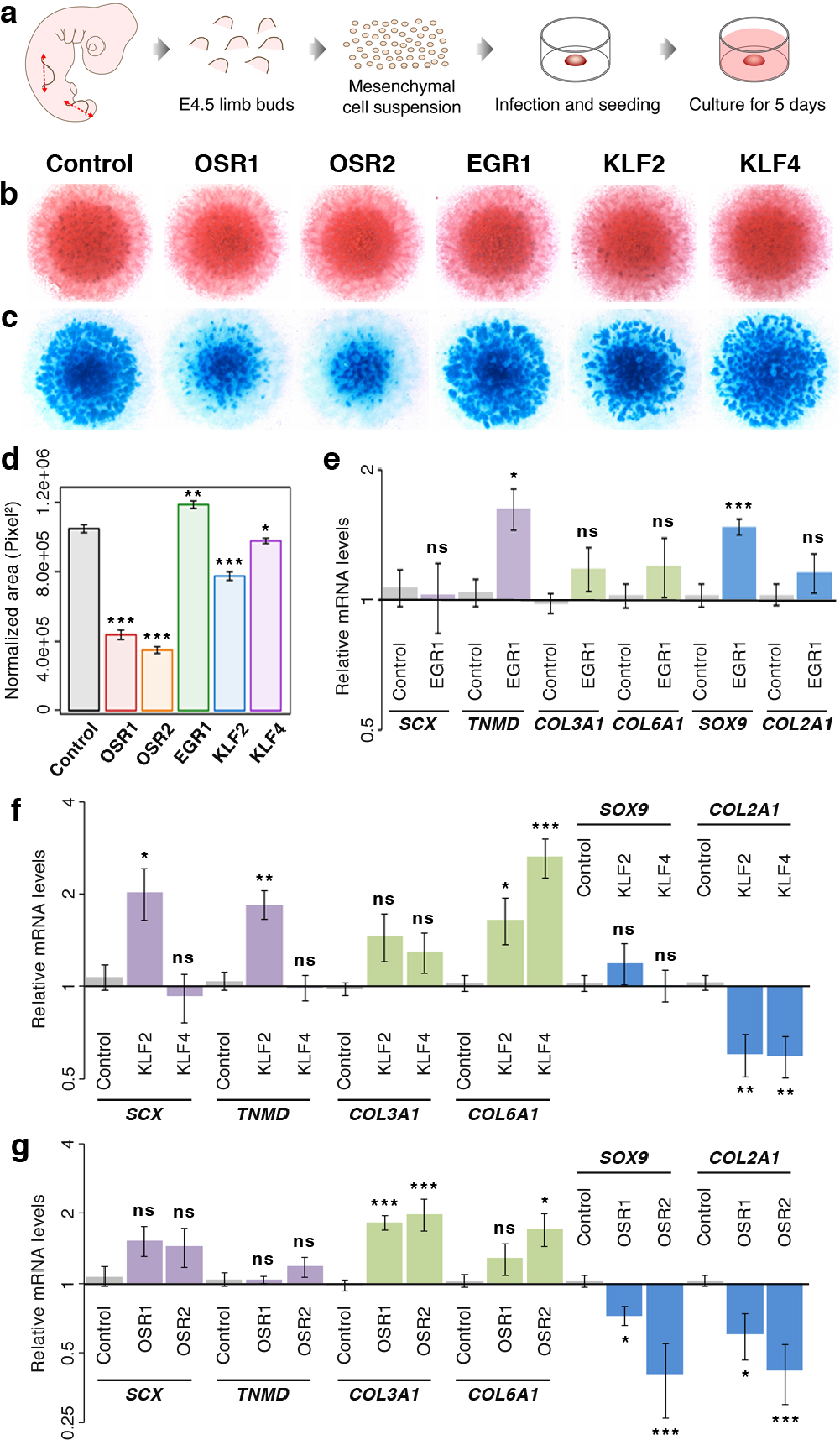
Differentiation of limb mesenchymal cells following TF overexpression. (**a**) Chick mesenchymal cells were isolated from E4.5 limb buds and cultured in high density for five days. (**b**) Eosin staining of TF-overexpressing chMM cultures. (**c**) Alcian blue staining of cartilage nodules formed in TF-overexpressing chMM cultures. (**d**) Quantification of chondrogenic matrix production: mean ±SEM; paired Student’s *t*-test: *, *P* < 0.05; **, *P* < 0.01; ***, *P* < 0.001. (**e**-**g**) Quantitative RT-PCR analysis of CT marker gene expression upon overexpression of EGR1 (**e**), KLF2/KLF4 (**f**) and OSR1/OSR2 (**g**) in chMM cultures. Graphs depict relative mRNA levels of *SCX* and *TNMD* (tendon markers), *COL3A1* and *COL6A1* (irregular CT markers), and *SOX9* and *COL2A1* (cartilage markers): mean ±SEM; two-tailed Mann-Whitney U test: ns, non-significant; *, *P* < 0.05; **, *P* < 0.01; ***, *P* < 0.001.

### Transcriptome analysis reveals similar regulatory functions between the five TFs

To gain insight into regulatory functions of the five TFs, transcriptome analysis was performed by RNA-seq. Total RNAs were extracted from two independent biological replicates of 5-day chMM cultures overexpressing each of the TF and subjected to high-throughput sequencing. Principal components analysis (PCA) and hierarchical clustering of the Euclidean distances on global gene expression profiles depicted a separation between the TF-overexpressing chMM cultures (Fig. 3a; Additional file 1: Fig. S1). Consistent with their similar expression domains in irregular CT, the gene expression profiles induced upon OSR1 and OSR2 overexpression were grouped together. In contrast, the gene expression profiles retrieved upon overexpression of the tendon-related TFs, EGR1, KLF2 and KLF4, were gathered together in a second group. However, the gene expression profiles upon KLF2 and KLF4 overexpression were more similar to each other than that induced upon EGR1 overexpression, which was consistent with their distinct expression domains associated with tendons. In summary, the gene expression profiles retrieved in the chMM cultures are in line with the limb expression patterns of the five TFs in the different CT subtypes.

**Fig. 3.**
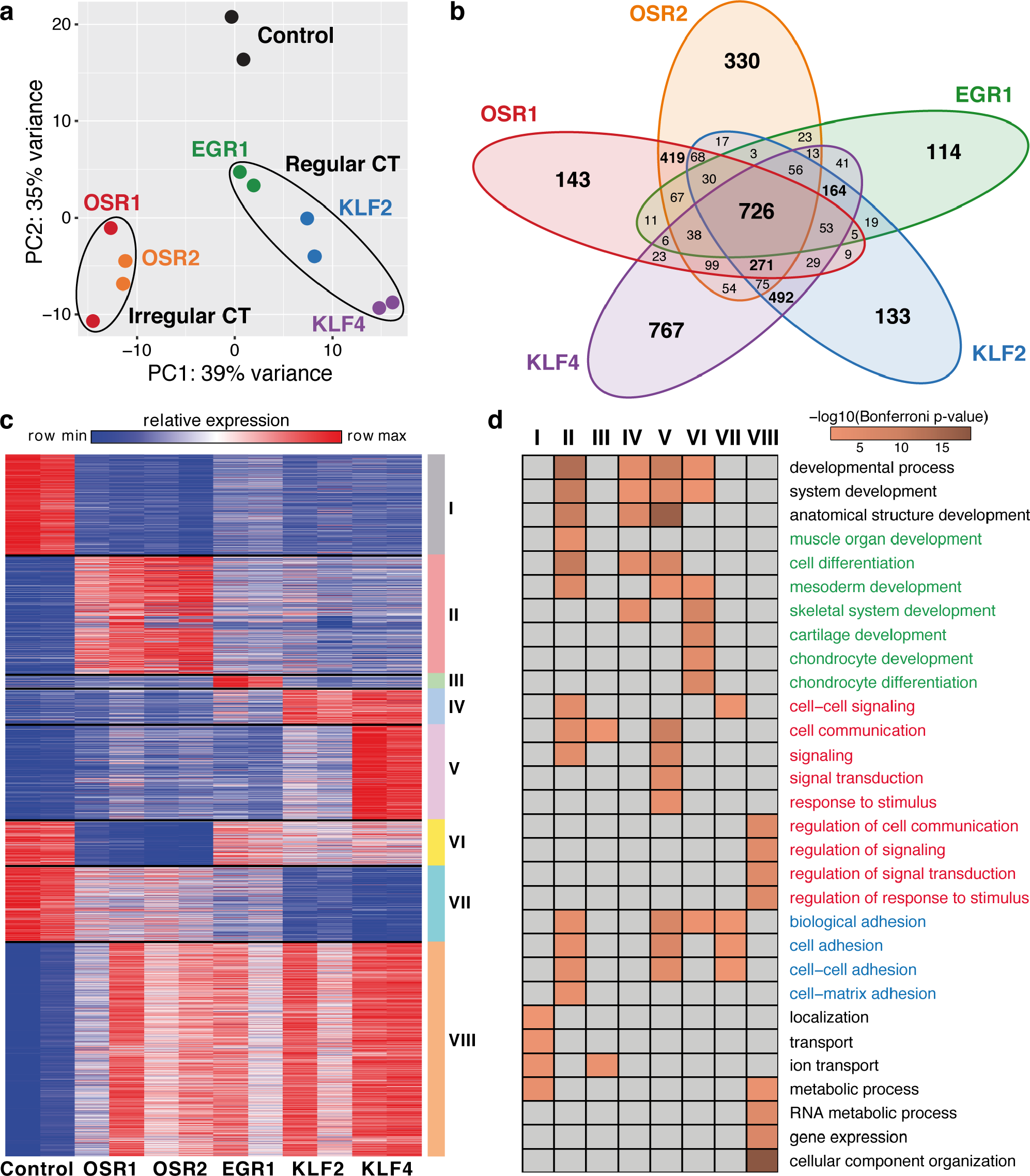
Gene expression profiles in chMM cultures upon overexpression of CT-associated TFs. (**a**) PCA analysis on global gene expression profiles of TF-overexpressing chMM cultures. (**b**) Venn diagram of the 4,298 non-redundant DE genes detected across all TF-overexpressing chMM cultures. (**c**) Gene clusters identified by *K*-means partitioning on the 4,298 non-redundant DE genes. (**d**) GO analysis for biological processes of the DE genes belonging to each *K*-means cluster. GO terms related to cell differentiation and development are depicted in green, cell signalling and communication in red, biological and cell adhesion in blue. Clusters having no significant enrichment for the specified GO terms are depicted in grey.

Following differential expression analysis, we identified between 1,369 and 2,907 differentially expressed (DE) genes for each TF-overexpressing culture compared to control cultures, resulting in a total of 10,712 DE genes for all TFs (Additional file 1: Fig. S2a-e). Interestingly, DE genes detected across all chMM culture conditions corresponded to a list of 4,298 non-redundant genes, indicating that the TFs shared common regulatory targets (Fig. 3b; Additional file 2: Table S1). While about one third (1,487; 34.6%) of DE genes were specific to a single TF, 2,811 (65.4%) DE genes were shared by at least two TFs (Fig. 3b). In addition, 726 (16.9%) DE genes were identified in all TF-overexpressing culture conditions (Fig. 3b). This indicates that the five TFs share a core of common regulatory targets, despite being expressed in distinct CT subtypes. In line with this, fold-change comparison (whether the gene is upregulated or downregulated) of the DE genes shared by at least two TFs revealed a high consistency among the TF regulatory patterns. Only 48 (1.7%) shared DE genes were identified as being regulated in opposite directions between the subset of TFs misregulating them (Additional file 1: Fig. S3). Therefore, the TFs did not have only similarities in the genes they regulated, but also in the manner these genes were affected.

Given the high consistency in the TF regulatory patterns and the elevated number of shared regulated genes, a gene clustering approach was performed on the 4,298 non-redundant DE genes by using *K*-means. This approach partitioned the DE genes into 8 clusters (Fig. 3c). A gene ontology (GO) analysis was then performed to identify potential biological processes enriched within each cluster (Fig. 3d). Genes downregulated by all five TFs were mainly involved in protein localization, ion transport and metabolic processes (Fig. 3c, d: cluster I). Genes upregulated by all five TFs were related to metabolic processes, gene expression, cellular component organization and several GO terms associated with the regulation of cell signalling and communication (Fig. 3c, d: cluster VIII). The clusters II, III, IV and V corresponded to genes upregulated specifically by one TF or by two closely related paralogous TFs (OSR1/OSR2 or KLF2/KLF4) (Fig. 3c). Genes in these clusters were mainly enriched for cell differentiation, mesoderm development, cell signalling/communication and biological/cell adhesion (Fig. 3d). Genes downregulated upon OSR1 and OSR2 overexpression were enriched for biological processes related to chondrogenesis (Fig. 3c, d: cluster VI), which was consistent with their anti-chondrogenic effect in chick cell cultures (Fig. 2c, d, g) [21]. Genes downregulated upon KLF2 and KLF4 overexpression were related to cell signalling and adhesion (Fig. 3c, d: cluster VII). In summary, the five CT-associated TFs differentially regulate the expression of genes mainly related to cell differentiation, signalling and adhesion. Thereby, they show a significant degree of overlapping regulatory function despite belonging to distinct CT subtypes.

### Molecular signatures downstream of the five TFs

The gene clustering approach revealed that a high proportion of genes upregulated by the selected CT-associated TFs was involved in signal transduction and biological adhesion. To further investigate this feature, signalling pathway enrichment analysis was performed on the complete set of DE genes identified for each TF. Of particular interest, signalling pathways related to extracellular matrix (ECM) components, such as integrin and cadherin signalling pathways, Wnt signalling, CCKR signalling and angiogenesis were enriched across all five TFs (Fig. 4a). Additional pathways were specifically enriched by a subset of TFs. For instance, TGF-β signalling pathway was identified upon overexpression of OSR1, OSR2 and KLF2, while Notch signalling pathway was enriched for both KLF2 and KLF4 DE genes (Fig. 4a). In addition, “axon guidance mediated by netrin” and “cytoskeletal regulation by Rho GTPase” pathways were enriched in both OSR1- and OSR2-associated DE genes (Fig. 4a). When comparing the averaged fold change across all TFs for each DE gene, it appeared that DE genes within each aforementioned pathway were significantly upregulated (Fig. 4b: median log2 fold change close to 1; Wilcoxon rank-sum test, *P* < 0.05). This tendency was not observed for the remaining non-DE genes associated with these signalling pathways (Fig. 4c: median log2 fold change close to 0). Nevertheless, a proportion of DE genes appeared rather downregulated for the integrin and TGF-β signalling pathways (Fig. 4b: lower whisker). This corresponded to a set of genes mainly repressed by both OSR factors (Additional file 1: Fig. S4a, b). Most of these genes encode collagens and BMP/GDF signalling molecules associated with cartilage and bone development, consistent with the anti-chondrogenic function of OSR1 and OSR2 (Fig 2c, d, g) [21]. In agreement with the signalling pathway enrichment analysis, overrepresentation test on the 4,298 non-redundant DE genes highlighted ECM, membrane and cytoskeleton cellular components (Fig. 4d). Altogether, gene expression profiling of chMM cultures overexpressing each TF supports a core of common regulatory functions across all TFs related to cell signalling, communication and adhesion. In addition, each TF (or paralogous TFs) also appear to be involved in e.g. regulation of individual signalling pathways, which could contribute to create a local microenvironment related to each CT subtype.

**Fig. 4.**
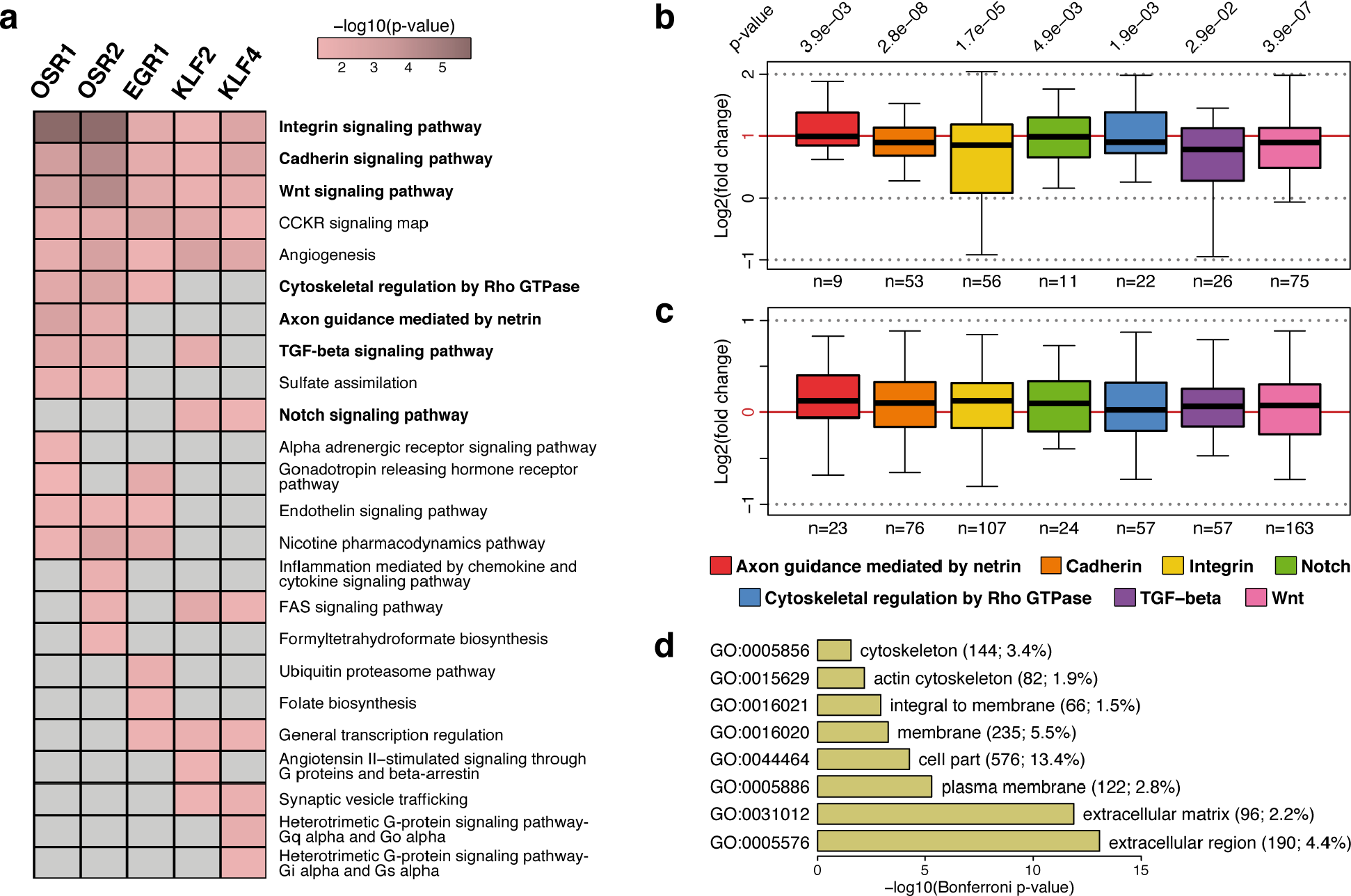
Signalling signature enrichment analysis of DE genes. (**a**) Panther pathways overrepresented within the DE genes detected upon overexpression of each TF in chMM cultures. DE genes having no enrichment for the specified Panther pathway are depicted in grey. (**b**, **c**) Global expression levels of DE genes (**b**) and non-DE genes (**c**) belonging to the selected Panther pathways. Log2 fold changes of each gene were averaged across all chMM culture conditions and replicates. Number of genes (n) in each Panther pathway is indicated at the bottom of each box. Paired Wilcoxon rank-sum test: p-values are indicated at the top of each box. (**d**) Cellular component GO analysis of the 4,298 non-redundant DE genes.

### Establishing genome-wide TF binding profiles

To further clarify the molecular mechanisms downstream of each TF, we investigated their genome-wide binding profile via ChIP-seq. Due to the absence of specific antibodies targeting each of the selected chicken TFs, we used the triple-FLAG (3F) tag that was fused C-terminally to the coding sequence (CDS) of each TF prior to their insertion into the RCAS-BP(A) vector. ChIP-seq was performed in two independent biological replicates of 5-day chMM cultures overexpressing each of the 3F-tagged TFs. Prior to ChIP, retroviral infection and overexpression of the recombinant TFs were assessed by immunohistochemistry and Western blot analysis against the 3F tag (Additional file 1: Fig. S5a, b). Similarity across all ChIP-seq signal profiles was assessed genome-widely in 500-bp non-overlapping windows by PCA. Comparison of the three first principal components partitioned the TF signal profiles by biological replicates and TF subgroups (Fig. 5a), indicating that paralogous TFs (OSR1/OSR2 and KLF2/KLF4) had a similar distribution across the genome.

**Fig. 5.**
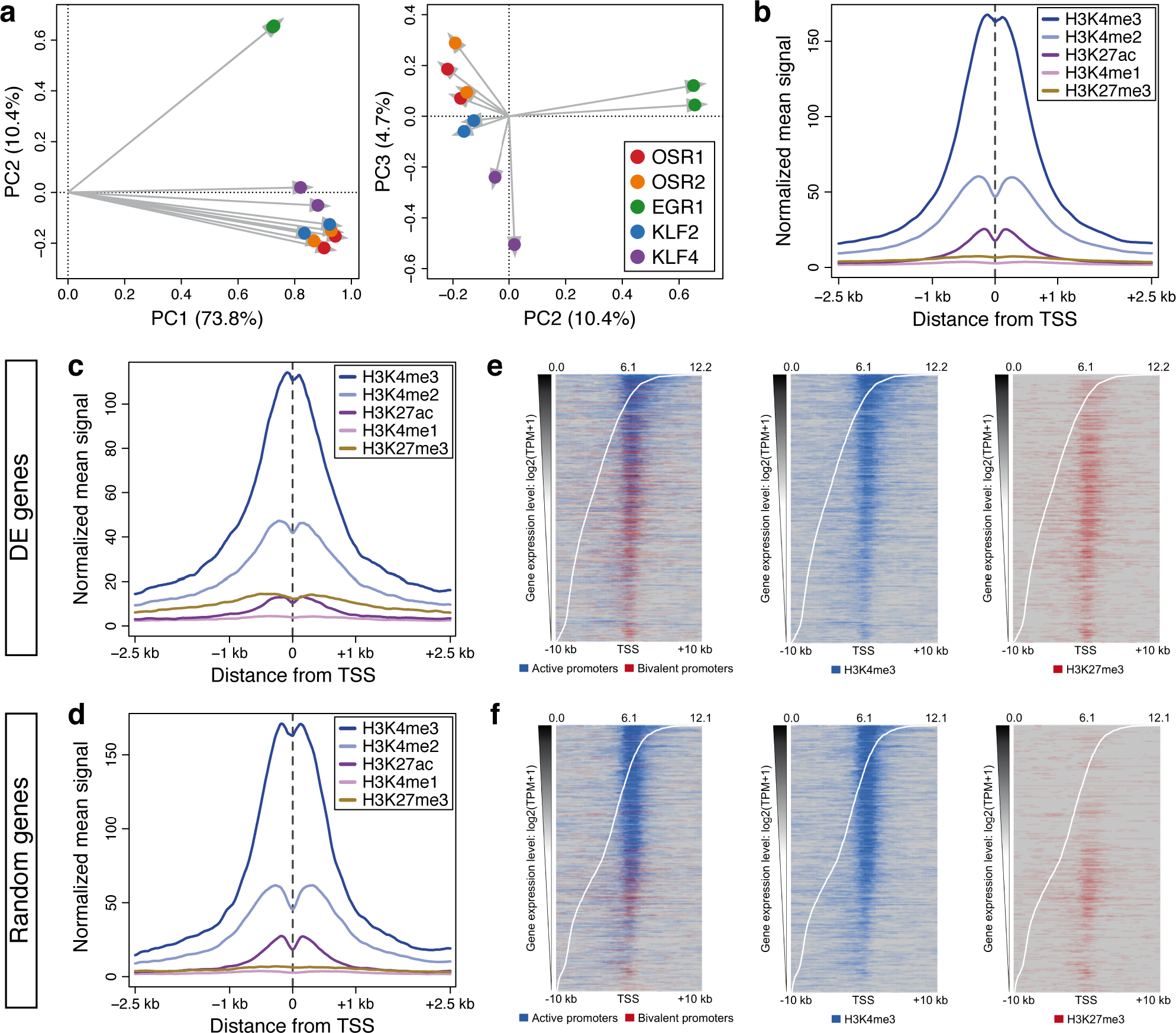
TF binding profiles and chromatin landscape in chMM cultures. (**a**) PCA analysis on the normalized ChIP-seq signal profiles of all TFs and biological replicates. (**b**-**d**) Normalized mean histone ChIP-seq signal surrounding the TSS of all genes (**b**), DE genes (**c**) and randomly selected genes (**d**). (**e**, **f**) Distribution of active (blue) and bivalent (red) promoter domains at the TSS of DE genes (**e**) and randomly selected genes (**f**). H3K4me3 (blue) signal is present in active and bivalent promoter domains, whereas H3K4me3 (red) signal is only detected in bivalent promoter domains. Intervals with a main regulatory domain being different from active and bivalent promoter are depicted in grey. Genes were ordered according to their expression levels (white curve).

Following peak calling, a total of 95,884 TF binding sites (TFBS) were identified (OSR1, 20,983; OSR2, 22,403; EGR1, 16,627; KLF2, 21,352; and KLF4, 14,519), which was 10-fold higher than the total number of DE genes. To potentially distinguish between functional and non-functional TF binding events, we decided to build a comprehensive map of the chMM chromatin landscape by characterizing promoter and enhancer regulatory domains. ChIP-seq experiments were performed against five histone post-translational modifications in two independent biological replicates of 5-day chMM cultures infected with retroviral particles overexpressing no recombinant protein, corresponding to the control conditions used for RNA-seq (Additional file 1: Fig. S6a). This approach was chosen to reveal the unbiased chromatin landscape in chMM cultures independently of TF overexpression. Mono-, bi- and tri-methylation of H3K4 (H3K4me1/2/3) were assessed to identify promoter and enhancer domains [23]. In addition, H3K27ac and H3K27me3 enrichments were investigated to distinguish between regions of transcriptional activity and facultative heterochromatin, respectively [23]. Based on this approach, we identified 20,427 promoters and 55,597 enhancers (Additional file 1: Fig. S6b, c). Surprisingly, we observed a decreased enrichment for histone marks associated with transcriptional activity (H3K4me3, H3K27ac) compensated by an increased signal of the repressive mark H3K27me3 at the transcriptional start site (TSS) positions of DE genes, as compared to their genome-wide levels (Fig. 5b, c: y-axis scale). This tendency was confirmed when investigating the TSS positions of a set of randomly selected genes of similar size (Fig. 5d: y-axis scale). This suggested the existence of bivalent promoter domains, which are known to be enriched in lineage-regulatory genes [24]. Consistently, DE genes were more associated with bivalent promoter domains at their TSS positions than randomly selected genes, regardless of their similar gene expression levels (Fig. 5e, f: left panels, white curve). By separating H3K4me3 (active and bivalent promoters) and H3K27me3 (bivalent promoters only) signals, it appeared that H3K4me3 active mark was overall less enriched at the TSS positions of DE genes than randomly selected genes (Fig. 5e, f: middle panels), whereas the H3K27me3 repressive mark displayed an opposite distribution (Fig. 5e, f: right panels). Although we cannot exclude that the increased ratio between repressive and active signal at promoters of genes affected by TF overexpression may reflect regulatory dynamics in the different cell populations [25], bivalent promoter domains suggest that DE genes are overall dynamically regulated and likely associated with CT differentiation and subtype-specific function, as opposed to housekeeping and ubiquitous genes that would be active and expressed across all cell types in limb cultures.

### Genome-wide patterns of TF occupancy

In order to assess TFBS functionality, binding locations identified for each TF were intersected with the regulatory domains. We focused on TFBS contained within promoters and enhancers, since we considered that binding events located in these regulatory domains likely contributed to the regulation of gene expression. Out of the 95,884 binding sites identified across all TFs, 31,289 (32.6%) overlapped promoter and enhancer regions, corresponding to 3,819-9,291 (17.9%-55.9%) binding events for each TF (Fig. 6a). De novo motif analysis was then performed on the 1,000 most significant binding sites for each TF. Recognition motifs identified for OSR1 and OSR2 were very similar and highly conserved with their known binding motifs in the fruit fly and the mouse (Fig. 6b) [26,27]. In agreement with previous reports, EGR1 and KLF4 binding motifs were enriched in cytosine/guanine (Fig. 6b) [26,28]. KLF2 recognition motif was highly consistent with the core binding sequence of the KLF protein family and was similar to the KLF4 secondary motif (Fig. 6b) [29]. Both binding motifs identified for KLF4 could contribute to its regulatory pattern observed in limb cell cultures, considering the 767 DE genes specifically identified for KLF4 and the 1,866 DE genes shared between KLF4 and KLF2 (Fig. 3b). Given the shared DE genes and the analogous recognition motifs, we then wondered whether the TFs depicted similarities regarding their occupancy within promoters and enhancers. Summit locations of all TFBS were retrieved and compared to each other. Binding events located within 500 bp surrounding peak summits (± 250 bp) were considered as shared binding locations. The 31,289 TFBS associated with promoters and enhancers corresponded to 17,714 binding regions. Out of these 17,714 regions, 10,979 (62.0%) were specific to a single TF, whereas 6,735 (38.0%) were shared by at least two TFs, including 1,045 (5.9%) that were common to the five TFs (Additional file 1: Fig. S7a). Shared occupancy among the TFs was further assessed by comparing the total number of shared binding sites that were identified between each TF pair. Consistent with their similar binding motifs, OSR1 and OSR2 shared more binding locations with each other than with any other TF (Additional file 1: Fig. S7b, c). Likewise, EGR1 and KLF4 tended to bind preferentially at similar regions as compared to the other TFs (Additional file 1: Fig. S7d, e). Surprisingly, we did not observe any preferential binding for KLF2 with any of the other TFs, including KLF4 (Additional file 1: Fig. S7f). Altogether, the binding motifs of the TFs and their genome-wide occupancy profiles appear consistent with the common and distinct regulatory patterns observed at the gene expression level.

**Fig. 6.**
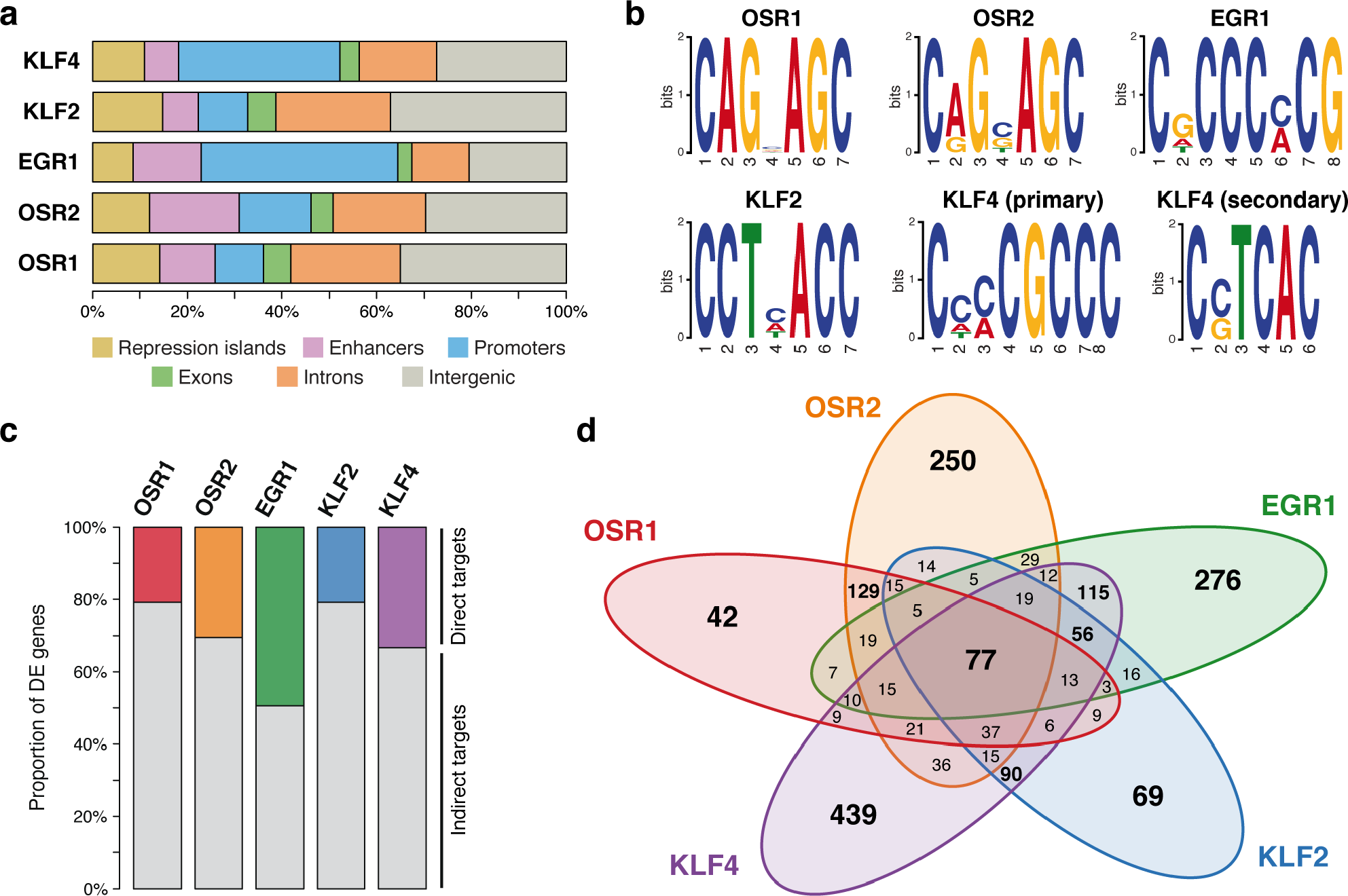
Regulatory patterns of CT-associated TFs. (**a**) Proportion of TFBS in chromatin domains and gene features. (**b**) TF recognition motifs. (**c**) Proportion of direct and indirect target genes of each TF in chMM cultures. (**d**) Venn diagram of the 1,858 non-redundant direct target genes detected across all TF-overexpressing chMM cultures.

### Identification of functional TFBS confirms a common regulatory core and distinct functions for the five TFs

To finally distinguish between indirectly and directly regulated genes, the 17,714 non-redundant binding regions identified across all TFs were intersected with the 4,298 DE genes identified from the RNA-seq data. Regions spanning from 10 kb upstream of the gene TSS to 10 kb downstream of the gene 3’-end were investigated for the presence of TFBS located within regulatory domains. This resulted in the identification of 3,210 genes that were potentially directly regulated by the TFs (Additional file 2: Table S1). The proportion of putative direct targets ranged from 20.9% (OSR1, 417; KLF2, 449) to 49.4% (EGR1, 677), depending on the TF (Fig. 6c). Consistent with the previous observations that the TFs shared common regulatory patterns and binding events in addition to their own specificity, the 3,210 genes considered as potential direct targets corresponded to 1,858 non-redundant genes (Fig. 6d). While 1,076 (57.9%) genes were directly regulated by a single TF, 782 (42.1%) genes were shared by at least two TFs, including 77 (4.1%) genes common to all TFs (Fig. 6d). In particular, OSR1 and OSR2 shared 318 target genes, while KLF4 possessed 317 target genes in common with EGR1 and 313 target genes in common with KLF2 (Fig. 6d). To further explore the regulatory link existing between the TFs, we next investigated whether the TFs sharing the same target genes tended to display similar binding locations. Consistent with the genome-wide occupancy analysis, OSR1 and OSR2 shared more binding locations with each other than with any other TF, albeit OSR2 displayed a higher binding specificity than OSR1 (Additional file 1: Fig. S8a, b). Likewise, EGR1 and KLF4 tended to bind preferentially at similar regions as compared to the other TFs, while also exhibiting their own binding specificity (Additional file 1: Fig. S8c, d). In contrast to the genome-wide occupancy analysis that did not depict any preferential binding between KLF2 and KLF4, we observed that both TFs tended to occupy similar locations in the vicinity of their common target genes, as compared to the other TFs (Additional file 1: Fig. S8e, f). In conclusion, the binding profiles of CT-associated TFs reflect their specificity and similarity in regards to their regulatory patterns.

### Validation of selected target genes

Coexpression of a TF and its putative target gene is a prerequisite for transcriptional regulation. Therefore, we compared the expression domains of TFs with that of selected candidate genes encoding signalling molecules in chick limbs. *NTN1* (netrin 1) was one of the genes that was upregulated in limb cell cultures upon overexpression of each TF (Fig. 7a; Additional file 1: Fig. S9a; Additional file 2: Table S1), through binding within an intronic enhancer (Fig. 7b). *NTN1* encodes a laminin-related secreted protein involved in axon guidance [30,31]. In E5.5 chick embryos, *NTN1* was expressed in both limb stylopod and zeugopod, displaying overlapping expression domains with those of all five TFs (Fig. 7c-h). At E8, *NTN1* was expressed in tendons, overlapping with *EGR1* expression domain close to muscle attachment (Fig. 7i, j). *NTN1*, *OSR1* and *OSR2* expression was observed in muscle CT at E9.5 (Fig. 7k-m). In addition, *NTN1* transcripts were detected in tissues delineating tendons at E9.5, similarly to *KLF2* and *KLF4* transcripts (Fig. 7n-p).

**Fig. 7.**
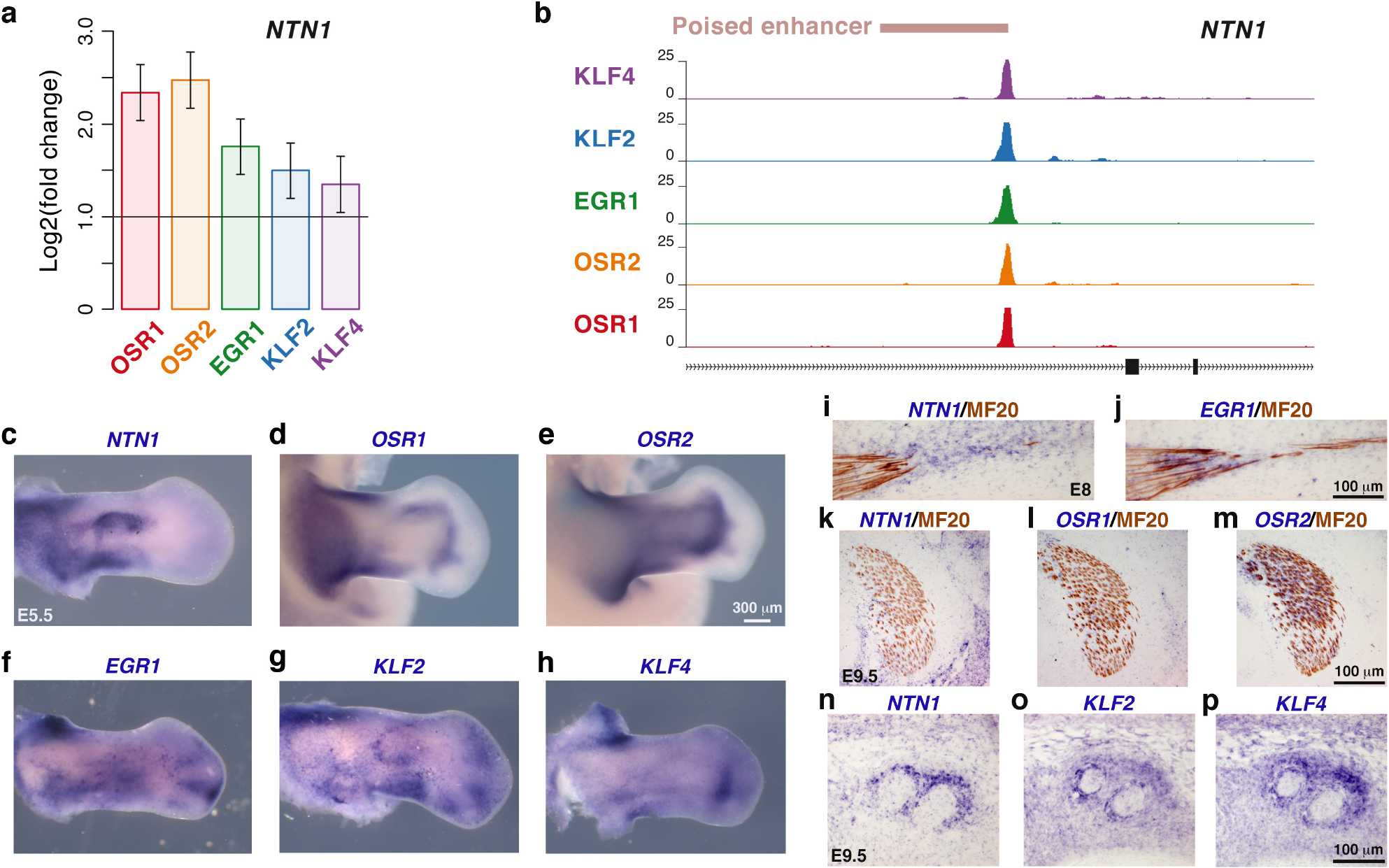
*NTN1* is a common target gene to the five CT-associated TFs. (**a**) *NTN1* expression levels in TF-overexpressing chMM cultures determined by RNA-seq. (**b**) Binding site within an intronic enhancer of *NTN1* gene for the five TFs identified by ChIP-seq. (**c**-**h**) Whole-mount in situ hybridization to hindlimbs of E5.5 chick embryos with *NTN1* (**c**), *OSR1* (**d**), *OSR2* (**e**), *EGR1* (**f**), *KLF2* (**g**) and *KLF4* (**h**) probes (blue). (**i**, **p**) In situ hybridization to forelimbs of E8 (**i**, **j**) and E9.5 (**k**-**p**) chick embryos followed by immunohistochemistry with the MF20 antibody (brown). Adjacent and longitudinal limb sections were hybridized with *NTN1* (**i**) and *EGR1* (**j**) probes (blue). Adjacent and transverse limb sections were hybridized with *NTN1* (**k**, **n**), *OSR1* (**l**), *OSR2* (**m**), *KLF2* (**o**) and *KLF4* (**p**) probes (blue).

Given the similar regulatory profiles of OSR1 and OSR2, we also selected *WNT11*, a common target gene of both TFs. *WNT11* encodes a secreted component of the non-canonical Wnt planar cell polarity pathway [32]. Both OSR factors increased *WNT11* expression in chMM cultures and bound at the same location within an intronic enhancer (Fig. 8a, b; Additional file 1: Fig. S9b; Additional file 2: Table S1). In E5.5 chick embryos, *WNT11* was expressed in limb mesenchyme, consistent with *OSR1* and *OSR2* expression patterns (Additional file 1: Fig. S9c-e). At E8 and E9.5, *WNT11* transcripts were detected in irregular CT within and surrounding muscles, overlapping with *OSR1* and *OSR2* transcripts (Fig. 8d-i). *WNT11* expression in CT downstream of OSR1 and OSR2 is consistent with the identified role of WNT11 in regulating muscle fibre type and orientation [33]. An additional target gene, *GDF6*, which encodes a secreted signalling factor of the TGF-β superfamily [34], was upregulated upon overexpression of OSR1 and OSR2 in chMM cultures (Fig. 8a; Additional file 1: Fig. S9b; Additional file 2: Table S1). However, only a binding site for OSR2 was detected in the vicinity of *GDF6*, indicating that OSR1 was not directly involved in the regulation of *GDF6* expression (Fig. 8c). In limbs of E5.5 and E8 chick embryos, *GDF6* expression domains overlapped with those of *OSR2* (Fig. 8j, k; Additional file 1: Fig. S9d, f).

**Fig. 8.**
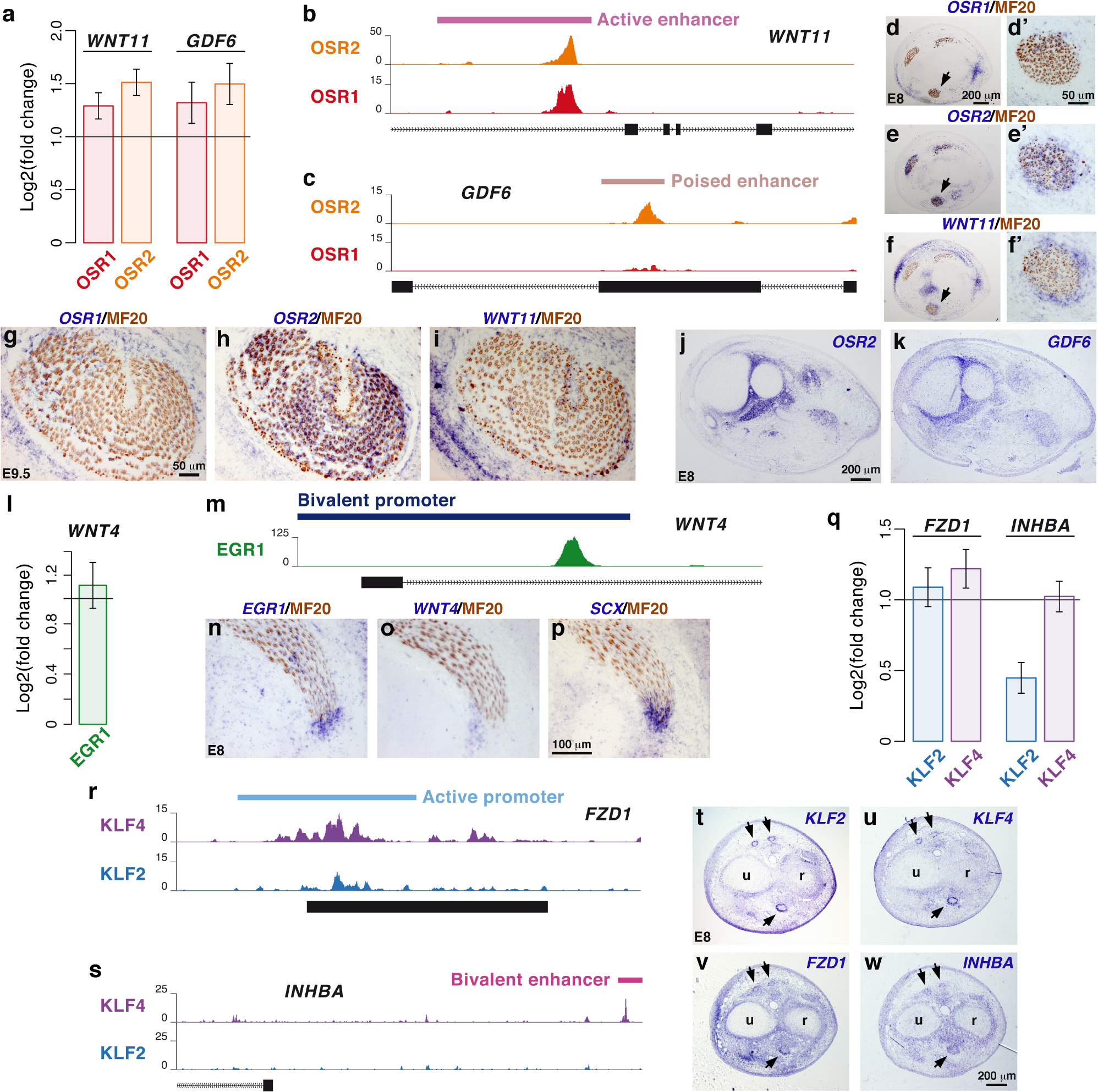
Selection of target genes encoding signalling molecules downstream of the five CT-associated TFs. (**a**) *WNT11* and *GDF6* expression levels in OSR1- and OSR2-overexpressing chMM cultures determined by RNA-seq. (**b**) Binding site within an intronic enhancer of *WNT11* gene for OSR1 and OSR2 identified by ChIP-seq. (**c**) Binding site within an exonic enhancer of *GDF6* gene for OSR2 identified by ChIP-seq. (**d**-**i**) In situ hybridization to forelimbs of E8 (**d**-**f**) and E9.5 (**g**-**i**) chick embryos followed by immunohistochemistry with the MF20 antibody (brown). Adjacent and transverse limb sections were hybridized with *OSR1* (**d**, **g**), *OSR2* (**e**, **h**) and *WNT11* (**f**, **i**) probes (blue). (**d’**, **e’**, **f’**) are high magnifications of the arrowed muscle in (**d**, **e**, **f**). (**j**, **k**) In situ hybridization to adjacent and transverse forelimb sections of E8 chick embryos with *OSR2* (**j**) and *GDF6* (**k**) probes (blue). (**l**) *WNT4* expression levels in EGR1-overexpressing chMM cultures determined by RNA-seq. (**m**) Binding site within the promoter of *WNT4* gene for EGR1 identified by ChIP-seq. (**n**-**p**) In situ hybridization to forelimbs of E8 chick embryos followed by immunohistochemistry with the MF20 antibody (brown). Adjacent and transverse limb sections were hybridized with *EGR1* (**n**), *WNT4* (**o**) and *SCX* (**p**) probes (blue). (**q**) *FZD1* and *INHBA* expression levels in KLF2- and KLF4-overexpressing chMM cultures determined by RNA-seq. (**r**) Binding site within the promoter of *FZD1* gene for KLF2 and KLF4 identified by ChIP-seq. (**s**) Binding site within an enhancer located upstream of *INHBA* gene for KLF4 identified by ChIP-seq. (**t w**) In situ hybridization to adjacent and transverse forelimb sections of E8 chick embryos with *KLF2* (**t**), *KLF4* (**u**), *FZD1* (**v**) and *INHBA* (**w**) probes (blue).

As a specific target gene of EGR1, we selected *WNT4*, which encodes a secreted member of the canonical Wnt signalling pathway [35]. *WNT4* was upregulated upon EGR1 overexpression in chMM cultures and was associated with an EGR1 binding site in its promoter region (Fig. 8l, m; Additional file 1: Fig. S9g; Additional file 2: Table S1). This is consistent with previous findings where EGR1 has been shown to bind upstream of *Wnt4* gene in the uterine endometrium during mouse pregnancy [36]. *EGR1* and *WNT4* displayed overlapping expression domains in proximal regions of forelimbs of E5.5 chick embryos (Additional file 1: Fig. S9h, i) and in tendons, close to muscle attachment in E8 limbs (Fig. 8n-p).

Given the common regulatory patterns of both KLF factors, we selected *FZD1*, which encodes a frizzled class receptor of Wnt signalling proteins [37]. *FZD1* was upregulated upon overexpression of KLF2 and KLF4 in chMM cultures (Fig. 8q; Additional file 1: Fig. S9j; Additional file 2: Table S1), and harboured a binding site for both KLF factors within its promoter region (Fig. 8r). In E8 chick embryos, *FZD1* was expressed in tissues delineating tendons, overlapping with the expression domains of *KLF2* and *KLF4* (Fig. 8t-v). Considering that KLF4 also displayed a distinct regulatory profile (Fig. 3c, 6d), we selected *INHBA*, which encodes the inhibin beta A subunit, a member of the TGF-β signalling pathway [38]. *INHBA* was upregulated upon KLF4 overexpression in chMM cultures (Fig. 8q; Additional file 1: Fig. S9j; Additional file 2: Table S1). In addition, a KLF4 binding site was located within an enhancer upstream of the TSS position of *INHBA* (Fig. 8s). In E5.5 and E8 chick limbs, *INHBA* and *KLF4* displayed overlapping expression domains in tissues delineating tendons (Fig. 8u, w; Additional file 1: Fig. S9k-m). Altogether, the selected target genes and their related CT-associated TFs exhibited overlapping expression domains in chick limbs.

### Common and divergent signalling/ECM signatures regulated by the CT-associated TFs

CT cells shape their microenvironment mainly by production of signalling/ECM molecules and/or via remodelling of the ECM. To explore this feature, we built a regulatory network on the 189 DE genes that were associated with seven of the previously identified signalling pathways (Fig. 4a, b; Additional file 2: Table S1). The resulting transcriptional network was composed of 513 interactions divided between 175 (34.1%) direct and 338 (65.9%) indirect connections (Fig. 9a). This network highlighted common and unique features for the CT-associated TFs. 38 (20.1%) genes were regulated by all five TFs, revealing a CT-typical signalling signature, whereas 47 (24.9%) genes were exclusively shared by paralogous TFs (OSR1/OSR2 and KLF2/KLF4) and 45 (23.8%) genes were specific to a single TF (Fig. 9a; Additional file 2: Table S1). The regulatory network was then subdivided for each individual TF to visualize the molecular interplay of each TF on selected signalling pathways (Additional file 1: Fig. S10-14). For instance, the Wnt signalling pathway was differently affected depending on the TF, with different sets of *WNT* ligand and *FZD* receptor genes regulated by each TF (Additional file 1: Fig. S10-14).

**Fig. 9.**
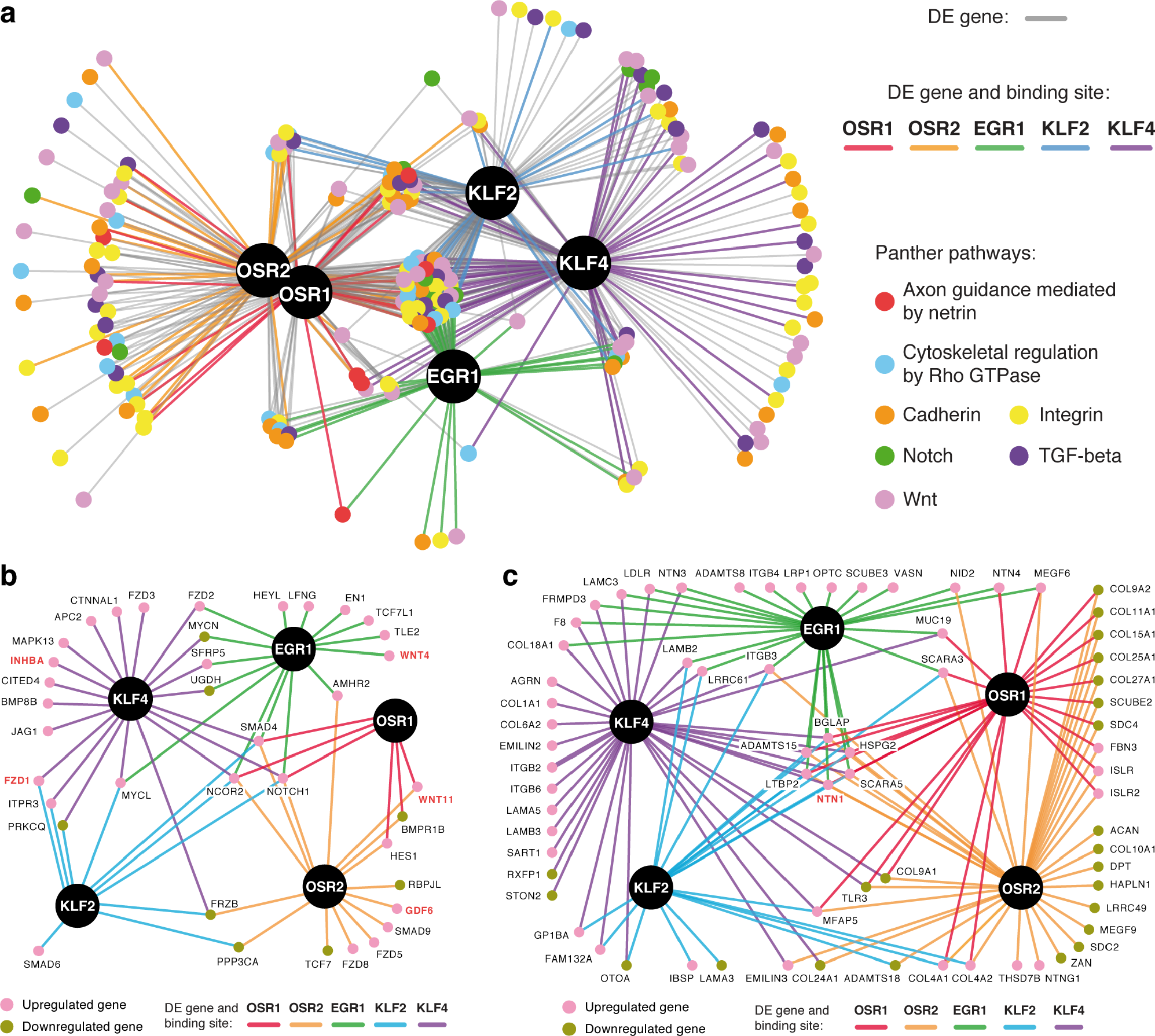
Regulatory networks of CT-associated TFs. (**a**) Transcriptional regulatory network of the CT-associated TFs and their target genes related to the indicated signalling pathways. Coloured connections correspond to direct interactions between the TFs and their target genes (DE gene and TFBS), while indirect interactions are depicted in grey (DE gene only). (**b**) Network representation of the 38 target genes associated with the Notch, TGF-β and Wnt signalling pathways that are directly regulated by the TFs. (**c**) Network representation of the 70 target genes associated with the ECM that are directly regulated by the TFs. Genes depicted in red correspond to the selected target genes investigated by in situ hybridization.

To reduce complexity, we then focused on the Notch, TGF-β and Wnt signalling pathways (Fig. 9b). Key components of these signalling pathways, *NOTCH1* (Notch receptor) and *SMAD4* (TGF-β signalling transducing protein) were regulated by all five TFs (Fig. 9b; Additional file 2: Table S1). By contrast, other genes were regulated by a subset of TFs. For instance, *WNT11* and *BMPR1B* (BMP receptor) were specific to OSR1 and OSR2, whereas *FZD1* and *PRKCQ* (protein kinase C theta) were regulated exclusively by both KLF factors (Fig. 9b; Additional file 2: Table S1). Lastly, we found genes that were specific to a single TF. This is the case for *GDF6* and *SMAD9* directly upregulated by OSR2, *WNT4* and *TCF7L1* (*TCF3*) directly upregulated by EGR1, and *INHBA* and *BMP8B* (BMP secreted ligand) directly upregulated by KLF4 (Fig. 9b; Additional file 2: Table S1). Altogether, this regulatory network identifies signalling genes that likely contribute to the biological function of all CTs or CT subtypes.

By finally focusing on direct target genes associated with the ECM, we found that the TFs regulated distinct, nevertheless partly overlapping molecular niches (Fig. 9c; Additional file 2: Table S1). *ADAMTS15* was directly upregulated by the five TFs and *ADAMTS8* was specific to EGR1, whereas *ADAMTS18* was directly downregulated by OSR2 and KLF2, (Fig. 9c; Additional file 2: Table S1). ADAMTS proteins are secreted metalloproteases with thrombospondin type-I motif that are involved in procollagen processing [39]. While ADAMTS15 and ADAMTS8 have proteoglycanolytic activity [39], mutations in *ADAMTS18* have been associated with bone disorders [40]. In addition, CT-associated TFs appeared to mediate collagen deposition by directly regulating genes encoding collagen α-chains. *COL4A1* and *COL4A2* were directly upregulated by OSR1, OSR2 and KLF2 (Fig. 9c; Additional file 2: Table S1). Type-IV collagen contributes to the assembly of basal lamina by binding to laminins [41]. By contrast, *COL9A1*, *COL9A2* and *COL11A1* were directly downregulated by OSR1 and OSR2 (Fig. 9c; Additional file 2: Table S1). Type-IX and XI collagens are known to form a network with type-II collagen in cartilaginous ECM [42]. KLF4 directly promoted the expression of *COL1A1* (Fig. 9c; Additional file 2: Table S1). Type-I collagen fibrils are the main component of tendons [7]. The ECM also acts as a source of developmental signals by sequestering and diffusing paracrine factors. The TFs appeared to directly mediate the positive expression of genes encoding laminin-related secreted netrins, such as *NTN1* (all five TFs), *NTN3* (EGR1 and KLF4) *NTN4* (OSR1 and EGR1) and *NTNG1* (OSR2) (Fig. 9c; Additional file 2: Table S1). In conclusion, the CT-associated TFs contribute to provide local patterning cues by mediating the expression of precise environmental molecules.

## Discussion

In this study, we have designed a transcriptional network downstream of five zinc finger TFs involved in CT differentiation. Two TFs (OSR1 and OSR2) were related to irregular CT, one TF (EGR1) to regular CT (tendon), and two TFs (KLF2 and KLF4) showed expression in CT delineating tendons. By combining different genome-wide strategies, we identified common and specific molecular signatures involved in limb CT differentiation. TF overexpression led to transcriptional activity changes in limb cells impacting on numerous cellular processes, including cell communication, migration and cell-cell/cell-matrix adhesion processes. Consistently, genes encoding signalling molecules, ECM components and cytoskeletal proteins appeared as regulated by the five TFs.

### Core molecular network downstream of the five TFs

We found 4,298 non-redundant DE genes upon overexpression of the five TFs. 65.4% of these genes were shared by at least two TFs, while 16.9% were common to all TFs. When direct regulation as judged by TF binding was considered, 77 genes were shared between the five TFs (Additional file 2: Table S1). We performed a conservative analysis and also restricted regulatory elements to a distance of 20 kb. It is well established that enhancer elements can be located further away, however identification of these regulatory interactions would include analysis of the 3D chromatin structure [43]. Altogether the number of common targets we identified is likely to be imperfect. Our data nevertheless show that the five TFs display common target genes despite their expression in different subcompartments of limb musculoskeletal system. This indicates that irrespective of CT type, whether it is specialized, dense regular or dense irregular, key molecular features are shared during the differentiation process of CT types, suggesting an archetypical CT signature.

One example for a common and directly regulated gene downstream of the five TFs is *NTN1*. Netrin 1 is a secreted ligand involved in axon guidance and developmental angiogenesis, in addition to preventing apoptosis triggered by one of its receptors, DCC (deleted in colorectal cancer) [44]. With the exception of EGR1, which only activates the expression of netrin ligands (*NTN1*, *NTN2* and *NTN3*), the other TFs positively regulate the expression of netrin receptors, *UNC5A* and/or *UNC5B* (Additional file 2: Table S1), known to mediate the netrin 1-induced axon chemorepulsion [44]. Our data suggests that NTN1 is an unexpected actor involved in migration and/or survival of CT cells during limb development. The molecular core downstream of the five TFs also includes a myofibroblast signature with *SRF*, *TAGLN* (*SM22*, transgelin), *TAGLN2* (transgelin 2), *CNN1* (calponin 1) and *ACTG2* (actin gamma 2) genes, which are positively activated by the five TFs, although not involving systematically direct binding sites (Additional file 2: Table S1). The myofibroblast signature is also confirmed by the presence of *SMAD4* in the list of the 77 common and directly regulated genes (Additional file 2: Table S1). *SMAD4* is a well-known profibrotic factor downstream of TGF-β1 [45]. The myofibroblast signature upon TF overexpression indicates that developmental CT differentiation shares molecular mechanisms with myofibroblast activation during fibrosis. In this context, it is noteworthy that *NOTCH1*, a component of a developmental signalling pathway described to be also involved in adult fibrosis [46], is a common and directly regulated gene downstream of the five TFs (Additional file 2: Table S1). The upregulation of Notch pathway components by each of the TFs suggests an unexpected involvement of the Notch signalling in limb CT formation during development.

### Specific regulatory patterns downstream of the five TFs

In addition to sharing a common molecular core, each TF displayed a specific regulatory pattern, albeit convergence was observed between related TFs, i.e. between OSR1 and OSR2 associated with irregular CT, and between EGR1, KLF2 and KLF4 mainly associated with regular CT. OSR1 and OSR2 are two markers of irregular CT and overexpression of each factor promotes the expression of irregular CT markers, such as *COL3A1*, while inhibiting that of cartilage markers in chick limb cells, as previously observed [21]. The list of OSR1 and OSR2 target genes identifies a molecular signature of irregular CT downstream of OSR factors in chick limb cells and potentially increases the limited number of irregular CT markers that are currently available. In addition, OSR1 and OSR2 represses the expression of genes related to cartilage lineage, which is consistent with the upregulation of cartilage-associated genes observed in *Osr1-*positive cells extracted from limbs of *Osr1* null mouse embryos [Vallecillo Garcia et al. in revision]. The master regulator of cartilage *SOX9* appears to be a direct target of OSR2, and the accessory regulatory factors *SOX5* and *SOX6* appear to be direct targets of both OSR1 and OSR2 in chMM cultures (Additional file 2: Table S1). OSR1 and OSR2 share 318 common target genes, but also display their own specificity. While only 42 target genes are unique to OSR1, 250 target genes are specifically regulated by OSR2 (Additional file 2: Table S1). The BMP ligand *GDF6* is one of the OSR2 specific target genes. GDF6 is known to play a role in establishing boundaries between skeletal elements during limb development, since inactivation of the *Gdf6* gene causes defects in joint, ligament and cartilage formation in mice [34]. This is in line with *Osr2*/*OSR2* expression in joint interzones in mouse [20] and chick embryos (Fig. 8j), as well as with the joint fusion defects observed in *Osr1*/*Osr2* double mouse mutants [47].

Although not being specific to tendons, EGR1 overexpression is sufficient to drive tendon cell differentiation in mouse mesenchymal stem cells [17]. Over 100 genes upregulated upon EGR1 overexpression were listed as being enriched in the transcriptome of *Scx*-positive cells isolated from limbs of mouse embryos [48]. This list includes *ADAMTS8*, *ADAMTS15*, *TAGLN*, *TAGLN2*, *FZD5* and *WNT4*, among others (Additional file 2: Table S1). *BMP4*, known to be expressed in chick limb tendons [49], was also positively regulated by EGR1 (although not directly) in our data (Additional file 2: Table S1). EGR1 is characterised as a fibrosis-promoting factor in many organs [50]. EGR1 has been also shown to directly regulate *Tgfb2* transcription in adult mouse tendons [17]. Here, we show that *TGFB1* (coding for the main fibrotic factor) is positively regulated by EGR1 (albeit not directly) in chick limb cells (Additional file 2: Table S1).

The exact function of KLF2 and KLF4 in limb musculoskeletal system formation is currently not known. However, we show here that both TFs display a striking expression delineating *SCX*-positive tendon/ligaments. In addition to the clear adhesion/migration signature downstream of both KLF factors in chick limb cells, KLF2 and KLF4 activates cell cycle genes such as *CDKN1A* (*P21*) and pluripotency genes (*SOX7*, *DKK1*, among others). Combined with the KLF2 and KLF4 well-known function in somatic cell reprogramming and pluripotency [51], this leads to the interesting idea that cells surrounding *SCX*-positive expression domains could be a source of tendon progenitors during development. Consistent with this idea, different tenogenic properties have been described for peritenon cells and tendon proper cells [52]. Beyond the 313 target genes that are shared between both KLFs and their similar binding occupancy in the vicinity of these target genes, KLF4 possesses 439 specific target genes. KLF4 specificity is corroborated with its primary binding motif that differs from the KLF core binding sequence identified in KLF2 and KLF4 ChIP-seq data. Indeed, the KLF4 binding site identified upstream of *INHBA* encompasses its primary recognition motif, whereas KLF2 and secondary KLF4 binding motifs are detected in the promoter region of their common target gene *FZD1*.

### A versatile molecular toolkit for shaping local niches

A significant proportion of directly or indirectly regulated DE genes comprises genes encoding signalling-associated molecules or ECM components and cell-matrix attachment molecules. The ECM is a three-dimensional insoluble network composed of secreted macromolecules, which provides positional and physical cues to influence cell position and migration [53,54]. Moreover, the ECM is a storage space for diverse growth factors that can be released upon e.g. proteolytic cleavage or mechanical stimulation. In this view, the ECM is both a scaffold structure and an integral part of cell-cell signalling mechanisms. Our data provide evidence that regional subspecification of limb bud mesenchymal tissues by TFs may be concomitant to local changes in the extracellular milieu. Importantly, we show that a common ECM signature activated by all five TFs exists for limb CT, while in addition each TF drives specific ECM and signalling factor genes. Individual TF or combinatorial TFs will impinge on the production of a particular ECM with specific growth factor decoration. This likely influences the behaviour of neighbouring tissues to create niches for invading cells. This is in line with the recognized importance of cell-ECM interactions for skeletal muscle, nerve and blood vessel development [55,56]. Thus, each TF or coordinated expression of a combination of TFs may be an elegant and adaptable way to achieve tissue (sub)compartmentalization in development.

## Conclusions

The transcriptional network presented here brings new insights into the molecular mechanisms orchestrating chick limb CT differentiation and function. The activity of CT-associated TFs impinges on overall common mechanisms including cell adhesion, migration and signalling. However, in addition to a common gene regulatory core downstream of the five TFs, each TF drives a specific program. These common and specific programs are likely to be at the root of tissue subspecification and local compartmentalization in developing limbs leading to the creation of local niches supporting organogenesis. This regulatory network offers valuable resources and opens new roads to better understand CT formation and function during limb development. In addition, such adaptable local transcriptional programs may apply to diverse contexts and might be a general principle of functional modulation.

## Methods

### Experimental procedures

#### Chick embryos

Fertilized eggs used for in situ hybridization were provided by the Institut de Sélection Animale (JA 57 strain, Lyon, France). Fertilized eggs used to prepare chMM cultures were obtained from VALO BioMedia (Lohmann Selected Leghorn strain, Osterholz-Scharmbeck, Germany). Embryos were staged according to the number of days in ovo at 37.5°C.

### Molecular cloning of the transcription factors

The CDSs of the chicken TFs OSR1, OSR2, EGR1, KLF2 and KLF4 were amplified by PCR by using the primers listed in (Additional file 3: Table S2). Cloning of the TF CDSs was performed by using a modified version of the pSlax-13 vector and the RCAS-BP(A) vector as previously described [57], with the exception that the 3F tag was fused C-terminally to each CDS.

### Chick micromass cultures

chMM cultures were prepared as previously described [58]. Briefly, limb buds were extracted from E4.5 chick embryos, ectoderm was dissociated by using a Dispase solution (Gibco) at 3 mg/mL, and limb mesenchyme was digested by using a solution composed of 0.1% Collagenase type Ia (Sigma-Aldrich), 0.1% Trypsin (Gibco) and 5% FBS (Biochrom) in 1X DPBS (Gibco). Prior to seeding, mesenchymal cells were mixed with retroviruses (1:1) and maintained in culture for 5 days at 37°C in DMEM/Ham’s F-12 (1:1) medium (Biochrom) supplemented with 10% FBS, 0.2% chicken serum (Sigma-Aldrich), 1% L-glutamine (Lonza) and 1% penicillin/streptomycin (Lonza). To assess cartilage differentiation, chMM cultures were fixed for 30 min with Kahle’s fixation solution (1% formalin, 30% ethanol and 4% acetic acid) and stained overnight at 4°C in 1% Alcian blue (Sigma-Aldrich) in 0.1 M HCl. Chondrogenic matrix areas were measured by using ImageJ [59]. For Eosin staining, chMM cultures were fixed overnight with 4% PFA in 1X PBS at 4°C and incubated for 2 min with 2.5 g/L of Eosin (Sigma-Aldrich) in 80% ethanol and 0.5% acetic acid. Viral 3F-tagged TF expression was monitored by using a mouse antibody directed against the 3F tag (Sigma-Aldrich, F1804; 1:500). Immunohistological staining was performed by using the Vectastain Elite ABC and the DAB Peroxidase Substrate kits (Vector Laboratories).

### RNA sequencing

Two biological replicates of chMM cultures were prepared from two independent pools of E4.5 limb buds and infected for 5 days with RCAS-BP(A) retroviruses overexpressing each of the TFs or no recombinant protein as control. For both replicates, RNA extracts were obtained by harvesting 6 chMM cultures with RLT buffer (Qiagen). Total RNAs were purified by using the RNeasy mini kit (Qiagen) in combination to a DNase I (Qiagen) treatment to prevent genomic DNA contamination. RNA libraries were prepared by using the TruSeq Stranded mRNA Library Preparation kit (Illumina), which enables to preserve the RNA strand orientation. Strand-specific 50-bp paired-end reads were generated by using a HiSeq 2500 sequencer (Illumina) with a mean insert size of 150 bp (Additional file 3: Table S3).

### ChIP sequencing

Harvesting of chMM cultures and ChIP experiments were performed as previously described [57]. Histone modification occupancy was investigated in two independent biological replicates of chMM cultures infected with RCAS-BP(A) retroviruses overexpressing no recombinant protein. 10 µg (8 chMM cultures) of chromatin extracts were incubated overnight at 4°C with gentle rocking with the following antibodies: 4 µg of mouse anti-H3K4me1 (Abcam, ab8895); 8 µL of mouse anti-H3K4me2 (Abcam, ab32356); 4 µL of mouse anti-H3K4me3 (Millipore, 07-473); 4 µg of mouse anti-H3K27ac (Abcam, ab4729); and 4 µg of mouse anti-H3K27me3 (Millipore, 07-449). TF binding profiles were investigated in two independent biological replicates of chMM cultures infected with RCAS-BP(A) retroviruses overexpressing each of the TFs. 30 µg (24 chMM cultures) of chromatin extracts were incubated overnight at 4°C with gentle rocking with 10 µg of mouse anti-FLAG (Sigma-Aldrich, F1804). Antibody-TF/Histone-DNA complexes were pulled down by using 40 µL of magnetic beads (Dynabeads protein G; Thermo Fischer). Ethanol-precipitated ChIP samples were resuspended in 46 µL of ddH20. Libraries were prepared by using the NEBNext Ultra DNA Library Preparation kit for Illumina (New England Biolabs). 50-bp single-end reads were generated by using a HiSeq 1500 sequencer (Illumina) (Additional file 3: Tables S4, 5). As input control, sonicated DNA from the nuclear fraction of each sample used for the ChIP procedures was also sequenced.

### In situ hybridization

Endogenous expression of the TFs was assessed by in situ hybridization on paraffin-embedded tissue sections. Chick embryo limbs were fixed overnight at 4°C in 60% ethanol, 30% formaldehyde at 37% and 10% acetic acid, and further processed as previously described [60]. For whole-mount in situ hybridization, chick embryos were fixed overnight at 4°C with 4% formaldehyde in 1X PBS and processed as previously described [61]. The following probes were used: *cOSR1* and *cOSR2* [20]; *cEGR1* [16]; *cKLF2* and *cKLF4* [62]. Expression of tendon and myogenic markers were assessed with the following probes: *cSCX* [15]; *cMYOD* [63]. Primers listed in (Additional file 3: Table S2) were used to generate probes detecting the following genes: *cFZD1*; *cGDF6*; *cINHBA*; *cNTN1* [64]; *cWNT4* (GEISHA ID, WNT4.UApcr); *cWNT11* (GEISHA ID, WNT11.UApcr).

### Quantitative RT-PCR analysis

Total RNAs were isolated from independent biological replicates of chMM cultures infected for 5 days with RCAS-BP(A) retroviruses overexpressing each of the TFs or no recombinant protein as control. RNA extracts were obtained as described for RNA sequencing. 500 ng of RNA extracts were used as template for cDNA synthesis using the High-Capacity cDNA Reverse Transcription kit (Applied Biosystems). Quantitative RT-PCR was performed by using the SYBR Green PCR Master mix (Applied Biosystems) in duplicates. Relative mRNA levels were calculated according to the 2-ΔΔCt method [65]. ΔCts were obtained from Ct normalized with chick *RPS17* (*S17*) and *GAPDH*. For each investigated gene, the mRNA levels of control chMM cultures were normalized to 1. Statistical analysis was performed by using Mann-Whitney U test with the GraphPad Prism V6 software. Primers used for quantitative RT-PCR are listed in (Additional file 3: Table S2).

### Computational analysis

#### Gene expression profiles

RNA-seq strand-specific read pairs were mapped against the chicken genome galGal4 [66] by using TopHat2 v0.14 [67] (parameters: -r 150; -N 3; –read-edit-dist 3; –library-type fr-firststrand; -i 50; -G) and the gene annotation model previously generated [68]. Alignment maps were split by strand by using SAMtools v1.2 [69] according to their FLAG field (strand plus: -f 128 -F 16, -f 80; strand minus: -f 144, -f 64 -F 16). Fragments (both reads of a pair) mapped on gene features were counted by using featureCounts v1.4.6-p3 [70] (parameters: -p; -s 2; –ignoreDup; -B; -R). Chimeric fragments aligned on different chromosomes were taken into consideration to overcome the gene fragmentation due to the location of gene parts on multiple chromosome contigs [68]. Fragment counts were then normalized by using DESeq2 v1.8.1 [71] and transcript abundances were calculated as transcripts per million (TPM) values according to the formula described in [72]. To evaluate the discrepancy among biological replicates and conditions, a regularized-logarithm (rlog) transformation was applied to normalized fragment counts followed by PCA analysis and hierarchical clustering of the Euclidean distances [71]. Differential expression analysis was finally carried out by using DESeq2 and a false-discovery rate (FDR; alpha) of 0.01. Genes with an absolute fold change of at least 2 and a Benjamini- Hochberg adjusted p-value (padj) below 0.01 were considered as being differentially expressed (Additional file 2: Table S1). Heat maps were generated by using the function heatmap.2 from the R package gplots. For given gene lists, rlog transformed fragment counts were used as input and hierarchical clustering was performed according to the one minus Pearson correlation.

### *K*-means gene clustering

*K*-means clustering was performed on the normalized fragment counts of the DE genes by using GENE-E [73] with a row distance metric set at 1 minus Pearson correlation and 2,000 iterations. The number of *K* clusters was defined at 8 because lower values did not separate distinct gene clusters and higher values subdivided meaningful gene clusters.

### Gene ontology analysis

GO analyses were performed for given gene lists by using the PANTHER statistical overrepresentation test r20160321 [74] and the Bonferroni correction for multiple testing. The following annotations were interrogated: PANTHER version 10.0 released on 2015-05-15 for GO-slim biological process, cellular component and pathways; GO ontology database released on 2016-04-23 for GO biological process complete.

### ChIP-seq coverage profiles

50-bp single-end reads generated for each ChIP and input fractions were first filtered on their quality by using the FASTX-Toolkit v0.0.13 [75]. Reads with a median quality value of minimum 28 were retrieved and mapped against the chicken genome galGal4 [66] by using BWA v0.5.9 [76] (default parameters). Uniquely mapped reads were then extracted and duplicated reads were finally removed by using the tool rmdup from SAMtools v1.2 [69]. Histone mark and TF coverage profiles were generated by using the tool bdgcmp from MACS2 v2.1.0.20140616 [77]. ChIP-seq signal was normalized independently for each biological replicate against the pooled input controls of both replicates according to the negative log10 of the Poisson p-value (-m ppois). Similarity between the TF binding profiles was assessed genome-widely in 500-bp non-overlapping windows by using PCA analysis with the R function prcomp (parameters: center TRUE; scale. TRUE).

### Histone modification peak calling

Peak calling for the histone ChIP-seq was performed as suggested by the ENCODE consortium [78]. For each histone modification, peaks were called independently for each biological replicate and for the pooled biological replicates, each time against the merged input control of both replicates, by using MACS2 v2.1.0.20140616 [77] (parameters: –bw 400, according to the sonicated DNA size;-g 1.0e9; –to-large). Except for the H3K27me3 mark, peak calling was performed twice for each replicate and pooled replicate: (i) narrow peaks passing a p-value (-p) of 0.01; and (ii) broad peaks passing an additional broad-peak p-value (-p 0.01; –broad; –broad-cutoff) of 0.1. Only broad peaks were called for the H3K27me3 ChIP-seq due to its diffused signal. Broad peaks detected for each replicate and pooled replicate that contain at least one narrow peak were extracted by using BEDtools intersect v2.24.0 [79]. Final sets of peaks for each histone modification were obtained by filtering broad peaks called for the pooled replicates that are shared between both biological replicates independently.

### Identification of regulatory domains

Regulatory domains were defined according to the combination of the different histone modification profiles obtained by ChIP-seq, independently of the gene annotation model and TSS positions given the fragmentation of the chicken genome [68]. Domains were divided into three categories: (1) promoters; (2) enhancers; and (3) repression islands. (1) Promoters were defined according to the presence of H3K4me3 signal. (2) Enhancers corresponded to regions enriched for H3K4me1 and devoid of H3K4me3 signal. (3) Repression islands were distinguished by the unique presence of H3K27me3 signal. Additional regions enriched for H3K4me2 but with no detectable H3K4me1 signal were classified as promoters, whereas regions containing both H3K4me1/2 marks were defined as enhancers. Promoter and enhancer domains were further subcategorised into four distinct states according to the active marks H3K4me3 and H3K27ac, and the repressive mark H3K27me3: (i) inactive, no active and repressive signal detected (H3K4me3^−^, H3K27ac^−^, H3K27me3^−^); (ii) poised, no active mark but repressive signal detected (H3K4me3^−^, H3K27ac^−^, H3K27me3^+^); (iii) active, only active mark detected (H3K4me3^+^ and/or H3K27ac^+^, H3K27me3^−^); and (iv) bivalent, both active and repressive marks detected (H3K4me3^+^ and/or H3K27ac^+^, H3K27me3^+^).

### Chromatin landscape at TSS positions

Normalized ChIP-seq signal was averaged for each histone modification from −2.5 to +2.5 kb surrounding the TSS of all genes, DE genes and randomly selected genes. To further investigate the increased enrichment of H3K27me3 mark, the 4,298 DE genes were filtered based on three criteria: (i) gene located on one single chromosome with a minimum size of 20 kb; (ii) gene body length of at least 1 kb; and (iii) −10/+10-kb regions around TSS within the chromosome borders. The resulting list was composed of 3,070 DE genes. The same criteria were applied to the randomly selected genes giving rise to a set of 3,080 random genes. 10-kb regions surrounding each TSS were retrieved and split into 100 intervals of 200 bp. For the genes having multiple transcripts with distinct TSS positions, the most upstream TSS was selected. Regulatory domains contained in each 200-bp interval were recovered in order to identify the most dominant domain per interval. Intervals marked with active and bivalent promoters were plotted in blue and red, respectively.

### Transcription factor peak calling

Quality of the TF ChIP-seq data was evaluated following the ENCODE consortium guidelines and metrics (Additional file 3: Table S5) [80]. Peaks were called by using MACS2 v2.1.0.20140616 [77] with low-stringency parameters to obtain a significant list of peaks (–bw 130/135, as determined by the cross-correlation analysis; -g 1.0e9; –to-large; -p 0.025). Irreproducible discovery rate (IDR) analysis was performed on the top 125,000 peaks according to their p-value [80] (parameters: peak.half.width −1; min.overlap.ratio 0; is.broadpeak F; ranking.measure p.value). The final set of TFBS was determined by selecting the number of peaks with an IDR threshold below 0.01 obtained from the pooled-replicate consistency analysis.

### Transcription factor occupancy

TFBS locations were intersected with regulatory domains and gene features by using BEDtools intersect v2.24.0 [79]. Summits of TFBS located within promoters and enhancers were retrieved and extended ± 250 bp. Extended summits were then merged by using BEDtools merge v2.24.0 [79]. Merged regions bound by at least two different TFs were further investigated to measure the genome-wide shared occupancy level between each TF pair. For each TF, the number of binding locations that intersected each of the four remaining TFs were counted separately and compared to its total number of shared binding locations. A similar approach was applied when analysing the shared occupancy level of the TFs in the vicinity of their target genes, albeit only the binding locations of the TFs directly regulating them were considered.

### Binding motif analysis

Motif analysis was performed by using DREME v4.11.2 [81] (default parameters) on the 150-bp sequences surrounding the summits (± 75 bp) of the 1,000 most significant TF peaks that overlapped with promoters and enhancers. Recognition motifs thus identified were then compared against motif databases by using Tomtom v4.11.2 [82] (default parameters).

### Transcriptional regulatory network

The regulatory network was built on the 189 DE genes that were regulated by at least one TF and associated with the selected signalling pathways using Cytoscape v3.4.0 and the edge-weighted spring-embedded layout [83]. The five TFs were determined as source nodes, while the DE genes were defined as target nodes. Interactions between each source node and its target nodes were marked as direct or indirect whether the differential expression was associated with a functional TFBS or not, respectively.

## Declarations

**Ethics approval and consent to participate:** not applicable.

**Consent for publication:** not applicable.

**Availability of data and material:** Sequencing data have been deposited on the Gene Expression Omnibus (GEO) database under the SuperSeries accession number GSE100517.

## Competing interests

The authors declare that they have no competing interests.

## Funding

This work was funded by the Deutsche Forschungsgemeinschaft (DFG; grant GK1631), the Université Franco-Allemande (UFA/DFH; grants CDFA-06-11 and CT-24-16), the Association Française contre les Myopathies (AFM; grants 16826 and 18626), the Fondation pour la Recherche Médicale (FRM; grant DEQ20140329500), the INSERM and the CNRS. MO was part of the MyoGrad International Research Training Group for Myology and received financial support from the FRM (grant FDT20150532272).

## Authors’ contributions

MO, DD and SS designed and conceived the study. MO, MM, GL, SN and MAB performed the experiments and collected the data. MO, MM, GL, SN, MAB, DD and SS analysed the data. STB and BT performed and supervised the RNA-seq procedure. JH performed and supervised the ChIP-seq procedure. MO, DD and SS interpreted the data. MO, DD and SS wrote the manuscript, with comments and approval from all authors.

## Acknowledgements

We are grateful to Stefan Mundlos (Charité Universitätsmedizin and Max Planck Institute for Molecular Genetics, Berlin, Germany) for generously sharing resources. We are thankful to the Sequencing Core Facility of the Max Planck Institute for Molecular Genetics for processing the RNA-seq. We are thankful to the Next Generation Sequencing Core Unit of the Berlin-Brandenburg Center for Regenerative Therapies for processing the ChIP-seq. We thank Peter Hansen and Peter N. Robinson (Charité Universitätsmedizin, BCRT, Berlin, Germany), as well as Marius van den Beek and Christophe Antoniewski (Institut de Biologie Paris-Seine, ARTbio, Paris, France) for providing access to Galaxy web servers. We thank Sophie Gournet (Institut de Biologie Paris-Seine, Paris, France) for illustrations.

## Additional files

**Additional file 1:** Supplementary figures S1-S14. (PDF)

**Additional file 2:** Supplementary table S1 combining RNA-seq and ChIP-seq data. (XLSX)

**Additional file 3:** Supplementary tables S2-S5. (XLSX)

## References

1. Heinz S, Romanoski CE, Benner C, Glass CK. The selection and function of cell type-specific enhancers. Nat Rev Mol Cell Biol. 2015;16:144–54.

2. Spitz F, Furlong EEM. Transcription factors: from enhancer binding to developmental control. Nat. Rev. Genet. 2012;13:613–26.

3. Nassari S, Duprez D, Fournier-Thibault C. Non-myogenic contribution to muscle development and homeostasis: the role of connective tissues. Front. Cell Dev. Biol. 2017;5:22.

4. Kalluri R. The biology and function of fibroblasts in cancer. Nat. Rev. Cancer. 2016;16:582–98.

5. Chevallier A, Kieny M, Mauger A. Limb-somite relationship: origin of the limb musculature. J. Embryol. Exp. Morphol. 1977;41:245–58.

6. Christ B, Jacob HJ, Jacob M. Experimental analysis of the origin of the wing musculature in avian embryos. Anat. Embryol. (Berl). 1977;150:171–86.

7. Gaut L, Duprez D. Tendon development and diseases. Wiley Interdiscip. Rev. Dev. Biol. 2016;5:5–23.

8. Kardon G. Muscle and tendon morphogenesis in the avian hind limb. Development. 1998;125:4019–32.

9. Lance-Jones C, Dias M. The influence of presumptive limb connective tissue on motoneuron axon guidance. Dev. Biol. 1991;143:93–110.

10. Michaud JL, Lapointe F, Le Douarin NM. The dorsoventral polarity of the presumptive limb is determined by signals produced by the somites and by the lateral somatopleure. Development. 1997;124:1453–63.

11. Kronenberg HM. Developmental regulation of the growth plate. Nature. 2003;423:332–6.

12. Braun T, Gautel M. Transcriptional mechanisms regulating skeletal muscle differentiation, growth and homeostasis. Nat. Rev. Mol. Cell Biol. 2011;12:349–61.

13. Huang AH, Lu HH, Schweitzer R. Molecular regulation of tendon cell fate during development. J. Orthop. Res. 2015;33:800–12.

14. Murchison ND, Price BA, Conner DA, Keene DR, Olson EN, Tabin CJ, et al. Regulation of tendon differentiation by scleraxis distinguishes force-transmitting tendons from muscle-anchoring tendons. Development. 2007;134:2697–708.

15. Schweitzer R, Chyung JH, Murtaugh LC, Brent AE, Rosen V, Olson EN, et al. Analysis of the tendon cell fate using Scleraxis, a specific marker for tendons and ligaments. Development. 2001;128:3855–66.

16. Lejard V, Blais F, Guerquin MJ, Bonnet A, Bonnin MA, Havis E, et al. EGR1 and EGR2 involvement in vertebrate tendon differentiation. J. Biol. Chem. 2011;286:5855–67.

17. Guerquin MJ, Charvet B, Nourissat G, Havis E, Ronsin O, Bonnin MA, et al. Transcription factor EGR1 directs tendon differentiation and promotes tendon repair. J. Clin. Invest. 2013;123:3564–76.

18. Hasson P, DeLaurier A, Bennett M, Grigorieva E, Naiche LA, Papaioannou VE, et al. Tbx4 and Tbx5 acting in connective tissue are required for limb muscle and tendon patterning. Dev. Cell. 2010;18:148–56.

19. Kardon G, Harfe BD, Tabin CJ. A Tcf4-positive mesodermal population provides a prepattern for vertebrate limb muscle patterning. Dev. Cell. 2003;5:937–44.

20. Stricker S, Brieske N, Haupt J, Mundlos S. Comparative expression pattern of Odd-skipped related genes Osr1 and Osr2 in chick embryonic development. Gene Expr. Patterns. 2006;6:826–34.

21. Stricker S, Mathia S, Haupt J, Seemann P, Meier J, Mundlos S. Odd-skipped related genes regulate differentiation of embryonic limb mesenchyme and bone marrow mesenchymal stromal cells. Stem Cells Dev. 2012;21:623–33.

22. Ahrens PB, Solursh M, Reiter RS, Singley CT. Position-related capacity for differentiation of limb mesenchyme in cell culture. Dev. Biol. 1979;69:436–50.

23. Kim TK, Shiekhattar R. Architectural and functional commonalities between enhancers and promoters. Cell. 2015;162:948–59.

24. Mikkelsen TS, Ku M, Jaffe DB, Issac B, Lieberman E, Giannoukos G, et al. Genome-wide maps of chromatin state in pluripotent and lineage-committed cells. Nature. 2007;448:553–60.

25. Hong SH, Rampalli S, Lee JB, McNicol J, Collins T, Draper JS, et al. Cell fate potential of human pluripotent stem cells is encoded by histone modifications. Cell Stem Cell. 2011;9:24–36.

26. Badis G, Berger MF, Philippakis AA, Talukder S, Gehrke AR, Jaeger SA, et al. Diversity and complexity in DNA recognition by transcription factors. Science. 2009;324:1720–3.

27. Meng X, Brodsky MH, Wolfe SA. A bacterial one-hybrid system for determining the DNA-binding specificity of transcription factors. Nat. Biotechnol. 2005;23:988–94.

28. Chen X, Xu H, Yuan P, Fang F, Huss M, Vega VB, et al. Integration of external signaling pathways with the core transcriptional network in embryonic stem cells. Cell. 2008;133:1106–17.

29. Sunadome K, Yamamoto T, Ebisuya M, Kondoh K, Sehara-Fujisawa A, Nishida E. ERK5 regulates muscle cell fusion through Klf transcription factors. Dev. Cell. 2011;20:192–205.

30. Serafini T, Colamarino SA, Leonardo ED, Wang H, Beddington R, Skarnes WC, et al. Netrin-1 is required for commissural axon guidance in the developing vertebrate nervous system. Cell. 1996;87:1001–14.

31. Dominici C, Moreno-Bravo JA, Puiggros SR, Rappeneau Q, Rama N, Vieugue P, et al. Floor-plate-derived netrin-1 is dispensable for commissural axon guidance. Nature. 2017;545:350–4.

32. Gao B. Wnt regulation of planar cell polarity (PCP). Curr. Top. Dev. Biol. 2012;101:263–95.

33. Gros J, Serralbo O, Marcelle C. WNT11 acts as a directional cue to organize the elongation of early muscle fibres. Nature. 2009;457:589–93.

34. Settle SH, Rountree RB, Sinha A, Thacker A, Higgins K, Kingsley DM. Multiple joint and skeletal patterning defects caused by single and double mutations in the mouse Gdf6 and Gdf5 genes. Dev. Biol. 2003;254:116–30.

35. DiRocco DP, Kobayashi A, Taketo MM, McMahon AP, Humphreys BD. Wnt4/β-catenin signaling in medullary kidney myofibroblasts. J. Am. Soc. Nephrol. 2013;24:1399–412.

36. Liang XH, Deng WB, Li M, Zhao ZA, Wang TS, Feng XH, et al. Egr1 protein acts downstream of estrogen-leukemia inhibitory factor (LIF)-STAT3 pathway and plays a role during implantation through targeting Wnt4. J. Biol. Chem. 2014;289:23534–45.

37. Laeremans H, Rensen SS, Ottenheijm HCJ, Smits JFM, Blankesteijn WM. Wnt/frizzled signalling modulates the migration and differentiation of immortalized cardiac fibroblasts. Cardiovasc. Res. 2010;87:514–23.

38. Howley BV, Hussey GS, Link LA, Howe PH. Translational regulation of inhibin βA by TGFβ via the RNA-binding protein hnRNP E1 enhances the invasiveness of epithelial-to-mesenchymal transitioned cells. Oncogene. 2016;35:1725–35.

39. Apte SS. A disintegrin-like and metalloprotease (reprolysin-type) with thrombospondin type 1 motif (ADAMTS) superfamily: functions and mechanisms. J. Biol. Chem. 2009;284:31493–7.

40. Wei J, Liu C, Li Z. ADAMTS-18: a metalloproteinase with multiple functions. Front. Biosci. (Landmark Ed). 2014;19:1456–67.

41. Mouw JK, Ou G, Weaver VM. Extracellular matrix assembly: a multiscale deconstruction. Nat. Rev. Mol. Cell Biol. 2014;15:771–85.

42. Fernandes RJ, Schmid TM, Eyre DR. Assembly of collagen types II IX and XI into nascent hetero-fibrils by a rat chondrocyte cell line. Eur. J. Biochem. 2003;270:3243–50.

43. de Laat W, Duboule D. Topology of mammalian developmental enhancers and their regulatory landscapes. Nature. 2013;502:499–506.

44. Cirulli V, Yebra M. Netrins: beyond the brain. Nat. Rev. Mol. Cell Biol. 2007;8:296–306.

45. Xu F, Liu C, Zhou D, Zhang L. TGF-β/SMAD pathway and its regulation in hepatic fibrosis. J. Histochem. Cytochem. 2016;64:157–67.

46. Hu B, Phan SH. Notch in fibrosis and as a target of anti-fibrotic therapy. Pharmacol. Res. 2016;108:57–64.

47. Gao Y, Lan Y, Liu H, Jiang R. The zinc finger transcription factors Osr1 and Osr2 control synovial joint formation. Dev. Biol. 2011;352:83–91.

48. Havis E, Bonnin MA, Olivera-Martinez I, Nazaret N, Ruggiu M, Weibel J, et al. Transcriptomic analysis of mouse limb tendon cells during development. Development. 2014;141:3683–96.

49. Wang H, Noulet F, Edom-Vovard F, Le Grand F, Duprez D. Bmp signaling at the tips of skeletal muscles regulates the number of fetal muscle progenitors and satellite cells during development. Dev. Cell. 2010;18:643–54.

50. Ghosh AK, Quaggin SE, Vaughan DE. Molecular basis of organ fibrosis: potential therapeutic approaches. Exp. Biol. Med. 2013;238:461–81.

51. Jiang J, Chan YS, Loh YH, Cai J, Tong GQ, Lim CA, et al. A core Klf circuitry regulates self-renewal of embryonic stem cells. Nat. Cell Biol. 2008;10:353–60.

52. Mienaltowski MJ, Adams SM, Birk DE. Tendon proper- and peritenon-derived progenitor cells have unique tenogenic properties. Stem Cell Res. Ther. 2014;5:86.

53. Charras G, Sahai E. Physical influences of the extracellular environment on cell migration. Nat. Rev. Mol. Cell Biol. 2014;15:813–24.

54. Rozario T, DeSimone DW. The extracellular matrix in development and morphogenesis: A dynamic view. Dev. Biol. 2010;341:126–40.

55. Eichmann A, Le Noble F, Autiero M, Carmeliet P. Guidance of vascular and neural network formation. Curr. Opin. Neurobiol. 2005;15:108–15.

56. Thorsteinsdóttir S, Deries M, Cachaço AS, Bajanca F. The extracellular matrix dimension of skeletal muscle development. Dev. Biol. 2011;354:191–207.

57. Ibrahim DM, Hansen P, Rödelsperger C, Stiege AC, Doelken SC, Horn D, et al. Distinct global shifts in genomic binding profiles of limb malformation-associated HOXD13 mutations. Genome Res. 2013;23:2091–102.

58. Solursh M, Ahrens PB, Reiter RS. A tissue culture analysis of the steps in limb chondrogenesis. In Vitro. 1978;14:51–61.

59. Schneider CA, Rasband WS, Eliceiri KW. NIH Image to ImageJ: 25 years of image analysis. Nat. Methods. 2012;9:671–5.

60. Wilkinson DG, Bailes JA, Champion JE, McMahon AP. A molecular analysis of mouse development from 8 to 10 days post coitum detects changes only in embryonic globin expression. Development. 1987;99:493–500.

61. Henrique D, Adam J, Myat A, Chitnis A, Lewis J, Ish-Horowicz D. Expression of a Delta homologue in prospective neurons in the chick. Nature. 1995;375:787–90.

62. Antin PB, Pier M, Sesepasara T, Yatskievych TA, Darnell DK. Embryonic expression of the chicken Krüppel-like (KLF) transcription factor gene family. Dev. Dyn. 2010;239:1879–87.

63. Pourquié O, Fan CM, Coltey M, Hirsinger E, Watanabe, Y Bréant C, et al. Lateral and axial signals involved in avian somite patterning: a role for BMP4. Cell. 1996;84:461–71.

64. Murakami S, Ohki-Hamazaki H, Watanabe K, Ikenaka K, Ono K. Netrin 1 provides a chemoattractive cue for the ventral migration of GnRH neurons in the chick forebrain. J. Comp. Neurol. 2010;518:2019–34.

65. Livak KJ, Schmittgen TD. Analysis of relative gene expression data using real-time quantitative PCR and the 2(-Delta Delta C(T)) Method. Methods. 2001;25:402–8.

66. Hillier LW, Miller W, Birney E, Warren W, Hardison RC, Ponting CP, et al. Sequence and comparative analysis of the chicken genome provide unique perspectives on vertebrate evolution. Nature. 2004;432:695–716.

67. Kim D, Pertea G, Trapnell C, Pimentel H, Kelley R, Salzberg SL. TopHat2: accurate alignment of transcriptomes in the presence of insertions, deletions and gene fusions. Genome Biol. 2013;14:R36.

68. Orgeur M, Martens M, Börno ST, Timmermann B, Duprez D, Stricker S. A dual transcript-discovery approach to improve the delimitation of gene features from RNA-seq data in the chicken model. bioRxiv. 2017; doi:https://doi.org/10.1101/156406.

69. Li H, Handsaker B, Wysoker A, Fennell T, Ruan J, Homer N, et al. The Sequence Alignment/Map format and SAMtools. Bioinformatics. 2009;25:2078–9.

70. Liao Y, Smyth GK, Shi W. featureCounts: an efficient general purpose program for assigning sequence reads to genomic features. Bioinformatics. 2014;30:923–30.

71. Love MI, Huber W, Anders S. Moderated estimation of fold change and dispersion for RNA-seq data with DESeq2. Genome Biol. 2014;15:550.

72. Wagner GP, Kin K, Lynch VJ. Measurement of mRNA abundance using RNA-seq data: RPKM measure is inconsistent among samples. Theory Biosci. 2012;131:281–5.

73. GENE-E. https://software.broadinstitute.org/GENE-E/. Accessed 28 Jun 2017.

74. Mi H, Dong Q, Muruganujan A, Gaudet P, Lewis S, Thomas PD. PANTHER version 7: improved phylogenetic trees, orthologs and collaboration with the Gene Ontology Consortium. Nucleic Acids Res. 2010;38:D204–10.

75. FASTX-Toolkit: FASTQ/A short-reads pre-processing tools. http://hannonlab.cshl.edu/fastx_toolkit/. Accessed 23 Dec 2016.

76. Li H, Durbin R. Fast and accurate short read alignment with Burrows-Wheeler transform. Bioinformatics. 2009;25:1754–60.

77. Zhang Y, Liu T, Meyer CA, Eeckhoute J, Johnson DS, Bernstein BE, et al. Model-based analysis of ChIP-Seq (MACS). Genome Biol. 2008;9:R137.

78. Kellis M, Wold B, Snyder MP, Bernstein BE, Kundaje A, Marinov GK, et al. Defining functional DNA elements in the human genome. Proc. Natl. Acad. Sci. U. S. A. 2014;111:6131–8.

79. Quinlan AR, Hall IM. BEDTools: a flexible suite of utilities for comparing genomic features. Bioinformatics. 2010;26:841–2.

80. Landt SG, Marinov GK, Kundaje A, Kheradpour P, Pauli F, Batzoglou S, et al. ChIP-seq guidelines and practices of the ENCODE and modENCODE consortia. Genome Res. 2012;22:1813–31.

81. Bailey TL. DREME: motif discovery in transcription factor ChIP-seq data. Bioinformatics. 2011;27:1653–9.

82. Gupta S, Stamatoyannopoulos JA, Bailey TL, Noble W, Maniatis T, Goodbourn S, et al. Quantifying similarity between motifs. Genome Biol. 2007;8:R24.

83. Shannon P, Markiel A, Ozier O, Baliga NS, Wang JT, Ramage D, et al. Cytoscape: a software environment for integrated models of biomolecular interaction networks. Genome Res. 2003;13:2498–504.

